# Investigation of sterile hydrogels as topical vehicles for APOSEC^TM^, a stressed peripheral blood mononuclear cell secretome for the treatment of poorly healing wounds

**DOI:** 10.64898/2026.02.26.708149

**Authors:** Dalia Hamid, Lisa Auer, Stefan Mohr, Sylwia Gazda-Miarecka, Melanie Salek, Hannes Kühtreiber, Nina Langoth-Fehringer, Tanja Pfleger, Victoria Klang, Michael Mildner, Clemens Aigner, Dirk Sorgenfrey, Hendrik Jan Ankersmit, Lea Ann Dailey, Gianluca Bello

## Abstract

APOSEC^TM^, a complex mixture of secreted proteins, lipids, and extracellular vesicles from stressed peripheral blood monocytes, is currently in clinical trials for the treatment of chronic, poorly healing wounds. When applied to open wounds, 1 mL reconstituted APOSEC^TM^ lyophilisate is syringe-mixed with 3 g sterile hydrogel prior to administration. This study investigates the pharmaceutical performance of this novel administration system. A gel formulation (APOgel) was developed for terminal sterilisation in pre-filled syringes with post-sterilisation viscosity (∼325-350□Pa*s at 1□s^-1^) comparable to a commercial benchmark gel. Syringe mixing of APOgel with a liquid APOSEC^TM^ surrogate (3:1) reduced viscosity by ∼67% but was highly reproducible across different operators (CV < 6%). Administration of three sequential dose units of the mixture from the syringe revealed an ∼20% higher content of active ingredients in the first and final dispensed compared to the middle unit, indicating non-uniform mixing in the closed syringe system. *In vitro* release studies over 72□h showed a 32% and 48% higher release of a small molecule marker and total proteins from the sterile APOgel compared to the benchmark gel as well as more pronounced gel swelling. However, efficacy studies in a murine wound healing model showed no significant difference between APOgel and the benchmark. These findings indicate that terminal sterilisation of gels for topical applications may provide benefits for more rapid release of active agents but syringe mixing of gels and a liquid requires optimisation to ensure uniform drug distribution.

**Highlights:**

- An autoclavable hydrogel for APOSEC^TM^ delivery was developed
- A novel syringe-mixing system for combining a gel with a liquid with subsequent dispensing of different volume units showed non-homogenous active ingredient distribution
- Final optimised APOSEC^TM^-APOgel formulation maintains functional wound-healing efficacy

## 1 Introduction

Chronic non-healing wounds represent one of the most common complications of diabetes and significantly increase the risk of amputation and premature death ^1^. The pathophysiology of these ulcers is multifactorial, involving impaired angiogenesis, excessive apoptosis of endothelial cells, and dysfunctional wound healing processes ^2^. Advanced therapeutic approaches, which range from cell-based to cell-free secretomes, skin substitutes and gene therapy, have shown promise in the treatment of such chronic, non-healing wounds ^1,2^. Amongst the therapies under development, the cell-free stressed peripheral blood mononuclear cell (PBMC) secretome (APOSEC^TM^) has shown promising tissue-regenerating activity in large and small animal wounding models ^3,4^. The mechanism of actions for the augmented wound healing associated with APOSEC^TM^ have been studied both in *in vitro* and *in vivo* systems ^5^. Beer et al. (2015) was the first to show that the bioactivity of APOSEC^TM^ correlated with the presence of a diverse array of secreted proteins, lipids, and exosomes ^6^. Studies indicated that stressing the PBMC with 60 Gy irradiation (either from a radioactive nuclide or linear accelerator) prior to cultivation in culture medium was indispensable for the APOSEC^TM^ activity ^3,6,7^. Bioinformatic analysis showed that the genes encoding the proteins released by stressed PBMC are mediators of regenerative pathways. Besides proteins, other key components include micro-RNA (miRNA), which influence cellular communication and regenerative processes ^6,8^. Evidence supports the claim that the wound healing effect is primarily caused by the initiation of angiogenesis and epithelial proliferation *in vitro* and *vivo*.

APOSEC^TM^ is produced under good manufacturing practice (GMP) conditions by γ-irradiation (60 Gy) of PBMCs extracted from donor blood followed by further culture in a proprietary medium (e.g. AM2 medium)^9^. Once the PBMCs are removed, the remaining liquid fraction, containing soluble factors and mediators (i.e., the secretome of the discarded PBMCs), is subjected to two orthogonal viral clearance steps and ultimately lyophilized (**Figure S1**; supplementary information section). Within this unique process, the culture medium becomes an integral component of the drug product. It is important to note that the cell-free APOSEC^TM^ is, by definition, a biological medicinal product (i.e. a culture-based product; Directive 2001/83/EC) as opposed to an advanced therapy medicinal product (ATMP), a gene therapy medicinal product, a somatic cell therapy medicinal product or a tissue-engineered product (Regulation EC No 1394/2007).

Validated biological assays have shown that APOSEC^TM^ can be produced with a compliant batch-to-batch consistency and a long lasting shelf-life stability^9–11^. The product successfully completed a phase I trial utilising autologous APOSEC^TM^ in 2017. In 2023, a phase I/II study for safety and clinical efficacy in the treatment of diabetic foot ulcers with allogeneic APOSEC^TM^ was finalised ^12,13^. During the trial, hospital pharmacy staff reconstituted the lyophilised APOSEC^TM^ (or the culture medium; placebo) with 0.2 mL 0.9% NaCl and aspirated this small volume into a syringe. A second syringe was manually filled with 0.8 mL Nu-Gel Hydrogel (Systagenix, Gatwick, West Sussex, UK), a sterile alginate-based gel for topical administration (Tables 1,2). Using a Luer connector (Combifix® adapter), the two syringes were connected and the liquid mixed with the gel, then collected in a single syringe. At the point of care, the mixture (∼ 1 mL) was applied directly to the open and a fresh wound dressing was applied (**Figure S2**; supporting information section). This procedure was repeated every 72 h.

Experiences during the trials highlighted two key areas where the administration system could be improved. First, it would be advantageous to have the sterile gel pre-filled into a syringe to negate the need for manually filling, thus reducing the preparation time and the risks of microbial contamination. Second, an increase in the amount of APOSEC^TM^ and gel per product could allow clinicians to flexibly administer 1, 2 or 3 mL depending on wound size (i.e. transition from a single dose to a multi-dose dispensing system). A study was designed to investigate the pharmaceutical performance of a putative system incorporating these two principles, referred to here as APOSEC 2.0 (**Figure 1**). In the first part of the study, we developed a hydrogel, similar in composition to the benchmark product, Nu-Gel, which could be pre-filled into an autoclavable cyclic olefin polymer (COP) syringe (3 g) and terminally sterilised. Ideally, the rheological properties of the autoclaved gel should be comparable Nu-Gel (∼350□Pa*s at 1□s^-1^) and no detectable degradation products should be observed.

**Figure 1:**
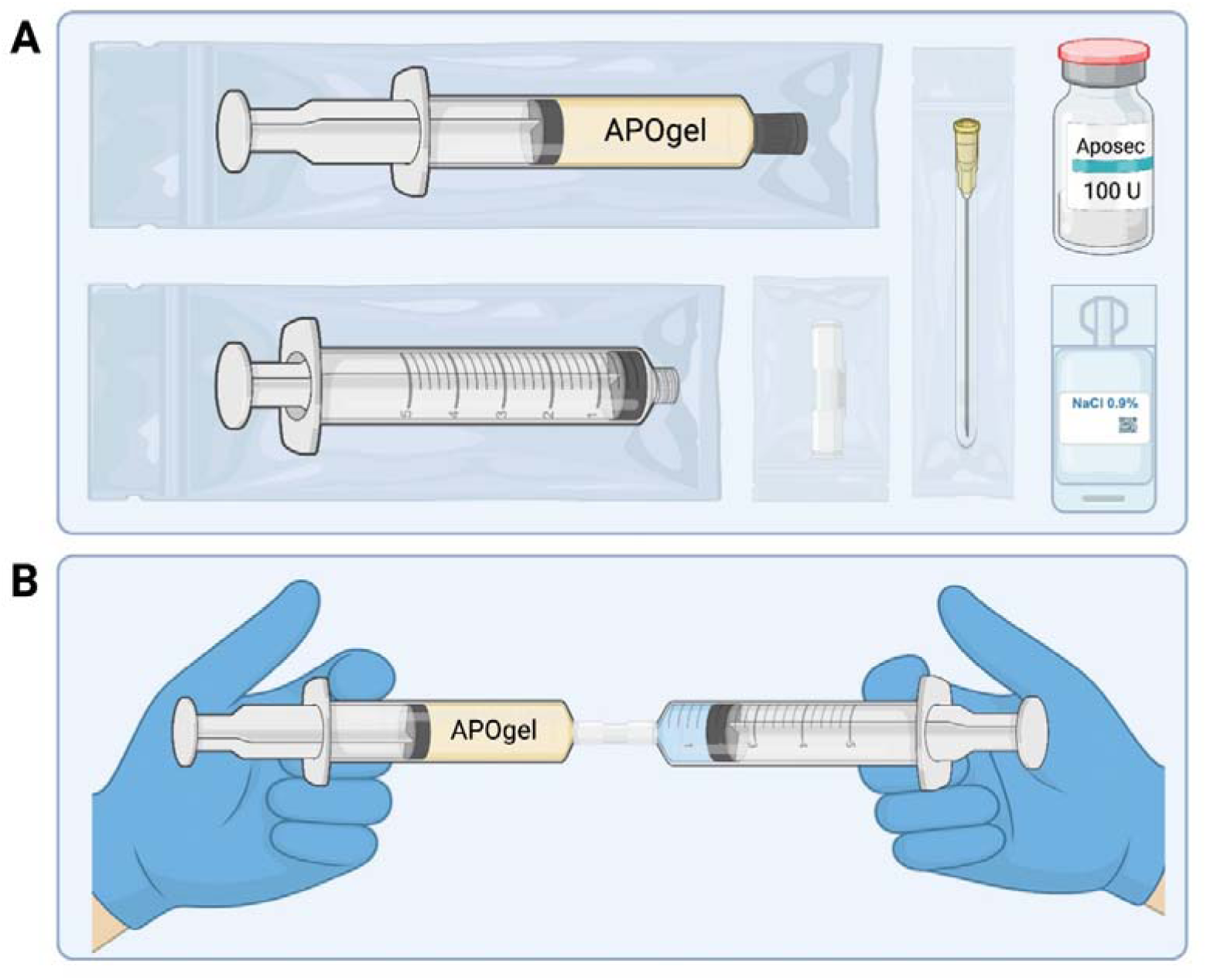
(A) Schematic of the APOSEC^TM^ product kit comprising a sterile bag containing a pre-filled COP syringe with 3 g sterile APOgel, a Luer connector, a vial of sterile, lyophilized APOSEC^TM^, and a vial containing 1 mL sterile isotonic saline for injections, a sterile, graduated syringe with needle for the resuspension of APOSEC^TM^, mixing with the gel and application to patients. (B) Illustration of the syringe-to-syringe mixing procedure of the reconstituted APOSEC^TM^ fluid using two Omnifix® Luer Lock solo syringes via Combifix® adapter.

In the second part of the study, the pharmaceutical performance of a syringe-mixing and multi-dose dispensing system were studied. The envisioned preparation procedure would involve 1) reconstitution of the lyophilised APOSEC^TM^ powder in 1 mL sterile saline solution, 2) aspiration of the liquid into a sterile, graduated 5 mL syringe, 3) connection of the liquid-filled syringe to the COP syringe containing 3 g sterile APOgel using a Luer connector (Combifix® adapter), 4) mixing of the liquid and gel by passing the contents back and forth between each syringe 20 times (**Figure 1B**), 5) collection of the mixture in the graduated syringe, and 6) administration of prescribed volume to the wound. The performance of such a dispensing system is not only interesting within the context of the APOSEC^TM^ development but also provides important insights for related drug delivery strategies, particularly in the emerging field of extracellular vesicle (EV)-based therapeutics. For example, there are an increasing number of studies investigating the co-formulation of therapeutic EVs together with gel-forming excipients for injectable in situ forming depot systems for localised, targeted delivery applications in tissue engineering or ocular drug delivery ^14–17^.

## 2 Materials and methods

### 2.1 Materials

Pharmaceutical grade carboxymethyl cellulose sodium salt Ph. Eur. (NaCMC), hydroxyethyl cellulose 250HX 10,000 mPa·s Ph. Eur. (HEC), propylene glycol PHE (PPG, Ph. Eur. 11.0) and sodium alginate Ph. Eur. 11.0 (NaAlg) were purchased from Gatt-Koller (Absam, Austria). Sodium chloride Ph. Eur. as ingredient of APOgel was from Fagron GmbH (Glinde, Germany). Sodium chloride (NaCl, purity ≥99 %) and casein soya bean digest (CASO) broth were provided by Carl Roth (Karlsruhe, Germany). A custom-formulated cell culture medium (AM2), containing Ringer’s lactate solution (838.75 mL/L),human serum albumin (8 g/L, or 0.120 mM) and recombinant human insulin (125 U/L, or 0.075 mM ^18^) for a ratio-weighted, total protein concentration of 0.195 mM, was purchased from PAN Biotech (Aidenbach, Germany) and was filtered through a 0.20 µm filter before use. Fluorescein sodium salt (SF) and phosphate-buffered saline (PBS) were obtained from Sigma Aldrich Ltd. (Dorset, UK). Water for injections (WFI) was produced in house according to the requirements of Ph. Eur. Monograph 0169 at the pharmacy of LKH-University Hospital Graz. Pierce™ Bradford Protein Assay Kit was purchased from Thermo Fisher Scientific (Massachusetts, USA). ClearJect® Luer Lock 5 mL cyclic olefin polymer (COP) syringes, suitable for autoclaving ^19^, were provided as a complete system with plunger stoppers, tip caps and plunger rods by Gerresheimer (Düsseldorf, Germany). AM2 medium was filled into Omnifix® Luer Lock Solo 5 mL 3-piece syringes, which were obtained, along with the corresponding Combifix® adapter (female to female), from B. Braun (Melsungen, Germany). Unjacketed Franz-type diffusion cells (orifice: 11.28 mm, receptor volume: 2 mL, with flange joint) were provided by PermeGear (Pennsylvania, USA), including magnetic stir bars and sealing rings. Hydrophilic Nuclepore™ polycarbonate membranes (pore size: 0.2 µm, diameter: 25 mm) were supplied by Cytiva (Massachusetts, USA). Parafilm® was purchased from Amcor Limited (Zürich, Switzerland). Nu-Gel was purchased from 3M (Minnesota, USA). Microorganism standards Vitroids™ of *Aspergillus brasiliensis* ATCC16404 and *Bacillus subtilis* ATCC6633 were purchased from Merck Millipore (Darmstadt, Germany).

### 2.2 APOgel production

A combination of factors, including intellectual property, sourcing, cost as well as manufacturing and scale-up considerations, contributed to the decision to employ a generic hydrogel system (APOgel), which could be produced under non-aseptic conditions (batch quantity: 2-3 kg), filled into COP syringes and terminally sterilised using steam heat sterilisation (121 °C, 2 bar, 15 min). The rheological properties of the sterilised APOgel should match the properties of the benchmark commercial product, Nu-Gel (3M), which is reported to have the composition ^20^ provided in **Table 1**. Since there are no literature reports on Nu-Gel excipient properties or the employed manufacturing and sterilisation methods, the proposed APOgel formulation was based loosely on the published Nu-Gel composition with selected modifications made during the product optimisation process.

**Table 1:** Reported composition of Nu-Gel ^20^ and the proposed composition of APOgel.

| Component | Nu-Gel composition<br>(% w/w) | APOgel composition<br>(% w/w) |
| --- | --- | --- |
| Water | 70.8 | 68.9 |
| Propylene glycol (PPG) | 25.0 | 25.0 |
| Sodium alginate | 3.0 | 3.0 |
| Carboxymethylcellulose (NaCMC) | 1.0 | 1.0 |
| Hydroxyethylcellulose (HEC) | 0.1 | 2.0 |
| NaCl | 0.1 | 0.1 |

It is well known that steam heat sterilisation leads to changes in viscosity and mechanical stability of hydrogels using cellulose ethers and alginate as gelling agents ^21^. Therefore, a higher initial viscosity of the APOgel formulation was required to account for the viscosity reduction observed during terminal sterilisation. Small scale pilot studies (50 – 250 g gel per batch) revealed that APOgel viscosity prior to terminal sterilisation needed to be > 700 Pa·s (at 1 s^-1^) to achieve a similar viscosity to Nu-Gel (∼400 Pas at 1 s^-1^) post-sterilisation (see electronic supplementary information; ESI). The increase in viscosity of the pre-sterilisation gel was achieved by increasing the amount of HEC in the formulation from 0.1% to 2% w/w (**Table 1**).

APOgel was produced in a class D cleanroom of the hospital pharmacy at LKH-University Hospital Graz under GMP conditions ^22^. Three replicate 3 kg batches were prepared. Briefly, the gelling agents were manually dispersed in PPG before being combined with NaCl dissolved in WFI. The ingredients were blended for 1.5 min at 750 rpm using a jacketed, vertical mixer (Stephan Mixer UM-12, ProXES GmbH, Hameln, Germany) under a vacuum of 0.8 bar below atmospheric pressure. This was followed by an additional 1 min mixing step at 1500 rpm. Finally, the set vacuum was maintained for 1 min without rotation. The gel was then filled into polypropylene wide-neck jars and stored at room temperature (20 - 25°C) until further use. For quality assurance reasons, all weights and process steps were documented in a manufacturing report.

Sterilisation of syringes containing APOgel was performed at a separate location under non-GMP conditions. APOgel samples were manually loaded into COP syringes (3.00 ± 0.03 g), then terminally sterilised using a vertical autoclave (Classic-Line HV-series from HMC Europe, Tüssling, Germany) set to the liquid program (121 °C, 2 bar, 15 min). After the autoclaving cycle, the syringes were allowed to cool to room temperature (23 °C, approximately one hour) followed by storage at 4 °C in darkness. Sterilised APOgel syringes were stored at 4 °C for a minimum of 24 h prior to further characterisation.

### 2.3 Rheological characterisation

The dynamic viscosity of all hydrogels investigated in the project was characterised using a modular compact rheometer (MCR 302, Anton Paar AG, Graz, Austria) with a 25 mm diameter cone-plate measuring system maintained at 23 °C via a Peltier heater. The gap distance between the stationary and the rotating plate was 0.1 mm. Zero-gap calibration was performed beforehand to ensure correct settings. A finite amount of hydrogel (400-600 mg) was dispensed onto the lower plate and trimmed after adjustment of the rotating plate, ensuring consistent gap filling. Dynamic viscosity was determined during increasing and then decreasing shear rates to assess linear rheological behaviour ^23^. Gels that were stored at 4°C were allowed to equilibrate to room temperature for approximately 1 h before measuring. The data was then analysed and compared through the Anton Paar RheoCompass software and plotted as viscosity (mPa*s) vs shear rates (s^-1^). The viscosity at selected shear rates of interest (1, 10 and 100 s^-1^) were also used for comparisons between different formulations.

### 2.4 Attenuated total reflection Fourier Transform Infrared spectroscopy (ATR-FTIR)

ATR-FTIR was used to characterise gel composition before and after sterilisation and compared to NU-Gel. NU-Gel and APOgel (5 g), before and after autoclave sterilization, were frozen in falcon tubes by plunging them into liquid nitrogen, followed by storage overnight (10-12 hours) at -80°C, to ensure complete freezing. The next day, the tubes were transferred unsealed (covering them with pricked parafilm to allow water and PPG to escape) into a lyophiliser to dry over three days. Spectra were obtained using an Alpha II FTIR-spectrometer (Bruker Optics GmbH & Co. KG, Ettlingen, Germany) equipped with an ATR module (Platinum-ATR, single reflection ATR with a monolithic diamond). Approximately 500 mg of dry gels were transferred directly onto the ATR crystal for measurements. At least three technical replicates, each consisting of at least 14 scans averaged and blank-corrected (air blank), were recorded with an optical resolution of 4 cm^-1^ between 4000 and 400 cm ^-1^. The processing of the data was carried out with SpectraGryph software (v1.2.16.1). The analysis of the transmittance (%T) spectra is carried out by comparing the spectra of APOgel, autoclaved APOgel and NU-Gel between each other and particularly, for the appearance of unusual peaks, related to the chemical structures of possible degradation products. To better identify such similarities and by-products, we focus on specific spectral regions common to the NaCMC, HEC and NaAlg components ^24,25^, here summarized in **Table 2**. Spectral similarities were assessed as described in the electronic supplementary information.

**Table 2:**
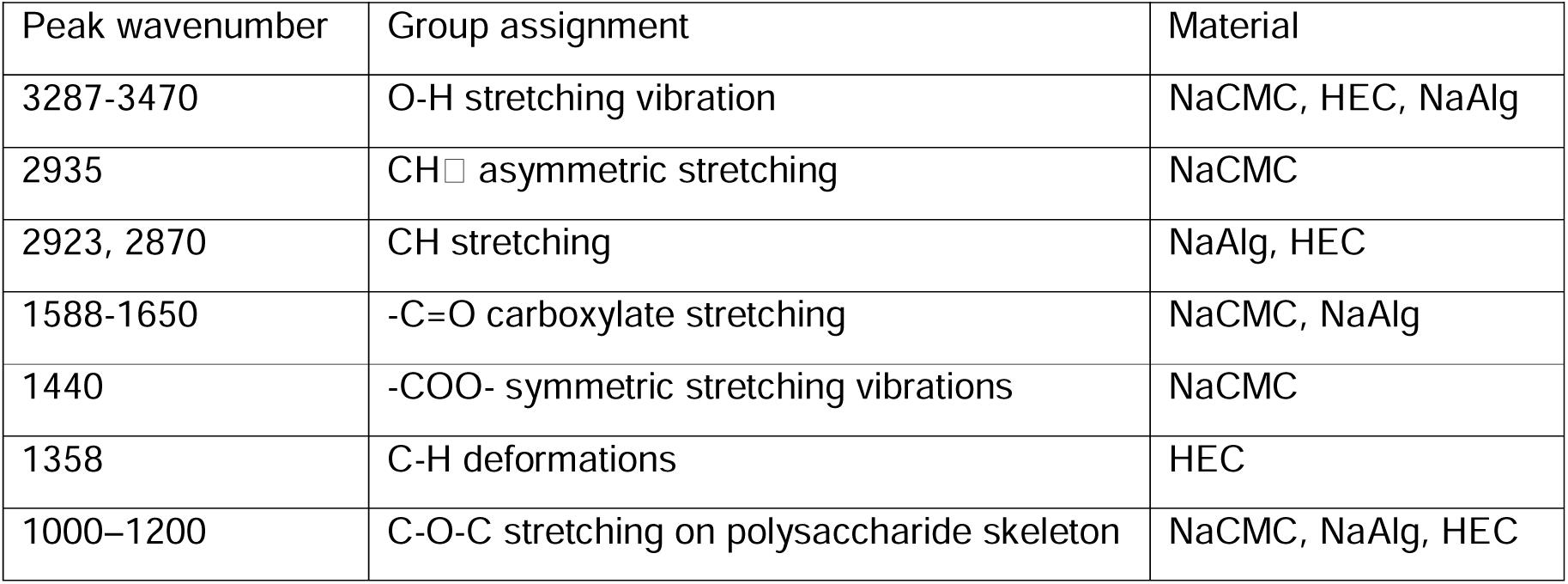
Group assignment from the ATR-FTIR spectrum for each dry components of the gel.

### 2.5 pH measurements

The relative pH of APOgel samples was determined before and after steam heat sterilisation. For measurement, samples were diluted to a final concentration of 10 mg/mL in distilled water (the pH of which was measured separately before dilution) to assess the influence of the gel on the pH of the water. Nu-Gel was used as a positive reference. The pH was measured using a calibrated Seven Compact pH meter (Mettler Toledo, USA) equipped with an InLab Micro sensor. pH values were recorded immediately after dilution, with n = 3 for each condition (APOgel before sterilisation, APOgel after sterilisation, Nu-Gel positive control, distilled water control).

### 2.6 Sterility test

The sterility test for topical use of a gel was carried out on two aerobic microorganisms following the directive of the European Pharmacopeia (Sterility 2.6.1) ^26^. The procedures were carried out under aseptic conditions in a BSL-2 safety cabinet, at a controlled temperature of 22.5 °C ±1°C and replicated three times (n=3). Briefly, sterile 12 mL glass tubes were filled with 10 g of casein soybean digest (CASO) broth and autoclaved. To assess the growth promotion suitability of the CASO broth for the aerobic microorganisms, the bacteria *Bacillus subtilis* ATCC6633 and the fungi *Aspergillus brasiliensis* ATCC16404 were inoculated at < 100 CFU. To evaluate the ability of APOgel to inhibit the growth of aerobes, 500 mg ± 1% of autoclaved APOgel were injected directly from the COP syringes into the vials, followed by inoculation of the respective bacteria at < 100 CFU (validation test). To determine whether the sterile CASO broth is suitable for the growth of the two strains, the media was inoculated with the microorganisms at < 100 CFU (growth promotion test). To determine the sterility of the autoclaved APOgel (sterility test), 500 mg ± 1% of gel were injected directly from the COP syringes into the vials. As the control, 500 mg ± 1% of distilled water autoclaved in COP syringes (with the same procedure as for APOgel) were injected directly into the vials. The growth promotion and validation tests were incubated for a maximum of three days for *Bacillus s.* and 5 days for *Aspergillus b*. The determination of the sterility of APOgel and the control was carried out after 15 days by visual analysis.

### 2.7 Characterisation of the syringe-mixing procedure, mixture viscosity, and content distribution

Pilot studies were used to establish an optimal procedure for combining the reconstituted APOSEC^TM^ liquid and APOgel using the closed-system syringe-mixing method described in **Figure 1**. To reduce costs, AM2 medium (the medium used to culture PBMCs during APOSEC^TM^ production) was used as a surrogate for the APOSEC^TM^ product. AM2 medium (1.00 ± 0.01 g) was filled into a sterile 5 mL graduated Omnifix® syringe and then connected to the COP syringe with 3 g sterile APOgel (or commercially available Nu-Gel, as a benchmark) via Luer adapter. The liquid was then transferred first to the COP syringe and subsequently the mixture was homogenised by transferring the contents back and forth between the two syringes for 20 total passes. The final mixture was then retained in the graduated syringe for dosing. The rheological behaviour of the mixtures was investigated as described in section 2.3 for all APOgel batches produced. To understand the effects of intra-operator variability on the viscosity of the mixture, three different volunteers were recruited to perform the mixing procedure after receiving the same verbal instructions and the rheological properties were characterised. Following this, the reproducibility of multi-dose dispensing was assessed by asking three volunteers to dispense three consecutive 1 mL doses from the syringes. The mass of each dispensed dose was measured gravimetrically.

The uniformity of APOSEC^TM^ components in each consecutive 1 mL dispensed dose was also assessed. Sodium fluorescein (SF; 0.025% w/w) was added to AM2 medium as an easily quantifiable surrogate for low molecular weight, water soluble APOSEC^TM^ components. Total proteins and peptides already present in the AM2 medium were evaluated using the Bradford assay. A 0.1 g sample was dispensed from the graduated syringe into a pre-weighed centrifuge tube and ∼ 1.5 g discarded (sample #1). This process was then repeated twice (samples #2-3). The mass of each sample was determined followed by dilution and quantification of SF and total proteins. SF content was quantified using fluorescence spectroscopy in a microplate reader (TecanTM infinite 200; Tecan Ltd., Männedorf, Switzerland) with excitation/ emission wavelengths of 485/ 535 nm. The AM2:gel mixture without the addition of SF was used for background subtraction. A calibration curve (0.0156-1 µg/mL; R^2^=0.993, LOD = 0.0052 μg/mL, LOQ = 0.0157 μg/mL) was used to quantify the amount of SF per g mixture. Total protein content was quantified using a Bradford assay according to manufacturer instructions ^27^. A standard curve of bovine serum albumin (2.5-25 µg/mL; R^2^=0.992, LOD = 1.158 μg/mL, LOQ = 3.509 μg/mL) was used to quantify the amount of total protein per g mixture. All experiments were performed in triplicate.

### 2.8 Release kinetics

Franz-type diffusion cells were used for *in vitro* release studies to evaluate the release profiles of SF and total protein from AM2:APOgel mixtures prepared by a single operator according to the procedure described in 2.7. Mixtures of AM2 and non-autoclaved Nu-Gel were prepared the same way and used as a benchmark control. A low-viscosity control was prepared containing a mixture of one-part AM2 medium combined with three parts APOgel formulation without gelling agents (i.e. NaCl, PPG and HPLC water). Diffusion cells were assembled and 2 mL PBS were added to the acceptor chamber under continuously stirring at 400 rpm. An infinite dose of each test sample (500 mg ± 100 mg) was placed onto a hydrophilic polycarbonate membrane (pore size: 0.2 µm, diameter: 25 mm) with a permeation area of 0.99 cm^2^. To reduce evaporation during the study, the sampling port was covered with aluminium foil and the donor chamber was occluded with Parafilm® and aluminium foil. All cells were placed in a thermostatically controlled shaking water bath (model 1092 from GFL; Burgwedel, Germany) at skin surface temperature (32±0.5 °C).

Two different study protocols were compared; 1) a 6-h study was performed to assess initial release plus diffusion rates under high gradient conditions and 2) a 72-h study was performed to assess release plus diffusion of SF or total proteins into the acceptor chamber over a therapeutically relevant timescale (i.e. a proposed dosing frequency of once every three days). In the 6-h study, samples (1 mL) were taken every hour for 6 h and replaced with an equal volume of PBS. For the 72-h release experiment, the entire volume of the receptor chamber (2 mL) was removed and replaced with PBS every 24 h. In the 72-h study, the entire Franz cell system was weighed at each time point to assessed evaporation losses and gel swelling.

SF and total protein were quantified as described in section 2.7. Release plus diffusion profiles were visualised by plotting the cumulative release as a percentage of the original amount of SF or total protein applied to the donor compartment against time (h). The cumulative release profiles for both the 6-h and 72-h studies were fitted using an exponential kinetic model developed for drug release from swollen gels under non-sink conditions ^28^. In general, sink conditions are maintained when drug concentration in the receptor medium does not exceed about 20% of its solubility, preventing saturation and allowing constant diffusion ^29^. In the current study, both the SF and proteins are freely soluble at all concentrations employed. Therefore, the 6-h release study is considered a high gradient condition, while the 72-h experiments reflect a more complex environment resulting from a reduction in the concentration gradient over time due to less frequent exchange of the receptor medium between samplings, as well as back-diffusion caused by the swelling gel. These conditions therefore demand kinetic modelling that goes beyond simple zero- or first-order models. To unify the fitting approach, the non-sink exponential release model described by Bernik et al (2006)^28^ (Equation 1) was applied for both time points.

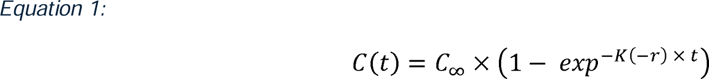

In Equation 1, the drug concentration (*C(t)*; %) at time *t* is proportional to the maximum predicted released concentration of drug at infinite time (*C*_∞_; %), which is expressed by the asymptote of the fitting curve. *C*_∞_ is reduced over time by the reabsorption, more precisely the back-diffusion, of drug by the swelling gel from the bulk, a process described by the reabsorption kinetic constant, *K(-r*). To understand the release kinetics from the gel to the bulk, *K(r),* at equilibrium we can calculate the kinetic constant of release by the equilibrium 2:

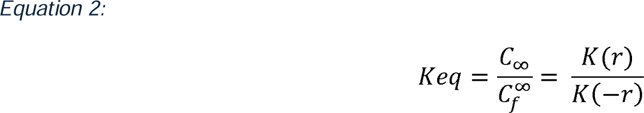

Where *C^∞^_f_* refers to the concentration of drug left in the gel at the equilibrium, a value that can be back-calculated from the experimental measurements.

### 2.9 *In vivo* model for comparative testing of APOSEC^TM^ hydrogel formulations

All animal experiments were approved by the Animal Ethics Committee of the Medical University of Vienna (GZ 2025-0.048.207) and conducted in accordance with national and institutional regulations for animal care and use. Female Balb/C mice (commonly used in wound healing studies) were obtained from Charles River Laboratories and housed under specific-pathogen-free conditions at the Core Facility for Biomedical Research of the Medical University of Vienna. Animals were maintained in individually ventilated cages under a 12 h light/dark cycle, with *ad libitum* access to standard food and water. After a minimum of a 14-day acclimatisation period, full-thickness excisional wounds (1 × 1 cm) were created on the dorsal skin under MMF anaesthesia (medetomidine-midazolam-fentanyl). Wound borders were outlined using a sterile 1 × 1 cm template and the marked skin area was carefully excised using surgical scissors. After the procedure, the anaesthesia was antagonized, and each mouse received a subcutaneous glucose solution for postoperative stabilization.

This study aimed solely to compare the *in vivo* equivalence of APOgel and Nu-Gel following mixing with APOSEC^TM^. For both hydrogels, APOSEC^TM^ lyophilisate (100 U; batch #732) was reconstituted in 0.5 mL 0.9% NaCl and mixed with the respective hydrogel (1.5 mL), consistent with the ratio described in section 2.7. From each final formulation, 0.5 mL were applied topically per mouse. The respective mixture was applied directly onto the wound surface (without wound covering) every second day for a total of ten days. Mice were monitored daily for general health and wound appearance. Standardized digital photographs of each wound were taken once per day, including a millimetre scale for calibration. Wound areas were quantified using ImageJ (Version: 2.16.0/1.54p) by manually tracing the wound margins and calculating the wound area in mm^2^ based on the calibrated scale. The percentage decrease in wound size was calculated relative to the day of maximum wound expansion, which served as the reference point for each animal. After ten days of observation, mice were euthanized by gradual CO_2_ inhalation.

### 2.10 Statistical analysis and graphics

Results were analysed using Microsoft Excel, GraphPad Prism (v. 10.01) and where specified, Python 3.13, for which the codes are provided. Wound areas were analysed using a two-way repeated-measures ANOVA with Geisser–Greenhouse correction (time = within-subject factor, treatment = between-subject factor). Images are produced using Microsoft Powerpoint and Biorender. The graphical abstract is created using icons from Freepik.com.

## 3 Results and discussion

### 3.1 APOgel characterisation

The quality requirements ^30,31^ for semi-sold drug products applied topically to open wounds is based on European Medicines Agency guidance ^32^ with the added specification of sterility, since the product will be applied to open wounds. Critical quality attributes (CQAs) defined for the sterile APOgel in the pre-filled syringe are listed in **Table 3**.

**Table 3:** CQAs and target specifications for the terminal sterilisation of APOgel in pre-filled COP syringes.

| Terminally sterilised APOgel pre-filled in COP syringes |  |  |
| --- | --- | --- |
| CQA | Test method | Specification |
| Sterility | Ph. Eur. Sterility 2.6.1 (for aerobic microorganisms) | No microbial growth of aerobic microorganisms detected |
| Viscosity | Dynamic viscosity | Post-sterilisation viscosity of APOgel within $\pm 15\%$ of Nu-Gel |
| Chemical stability | ATR-FTIR | No appearance of degradation peaks upon autoclaving |
| pH | pH-meter | pH within $\pm 15\%$ of Nu-Gel |
| Appearance | Visual inspection | Minimal changes in bubbles and colour compared to non-sterile gel |

Steam heat sterilisation is the most common method for terminal sterilisation of medicinal products and is recommended by regulatory agencies (European Pharmacopoeia 5.1.5, 2017, European Medicines Agency (EMA), 2019), as long as the product can withstand the elevated temperatures employed (121 °C, 2 bar, 15-20 min) ^21,33^. In the context of APOgel production as topical product, sterility can be validated for aerobic microorganisms following the European Pharmacopeia 2.6.1 ^26^. Sterility investigations showed that autoclaved APOgel has no inhibitory activity (validation test) towards *Aspergillus b.* but inhibits the growth of *Bacillus s*. Interestingly, after 5 days (beyond the validation test period set by the European Pharmacopeia) *Bacillus s.* grew in CASO broth in the presence of APOgel. The sterility tests carried out on sterile APOgel injected directly from the COP syringes showed no microbial growth after 15 days, moreover, the control validated the test. Hence, steam sterilisation in COP syringes is an effective method to sterilise APOgel.

Despite the advantages of steam heat sterilisation (efficiency, speed, simplicity, low cost and no residues in the product ^34,35^), the high temperatures needed can affect the structural properties of gel-forming polymers and can lead to the reduction of water content in the hydrogel due to evaporation ^36^. Additionally, water vapour can cause degradation and hydrolysis of some polymers, altering their gel structure and properties, especially if bubbles are present in the gel ^34,37^. This effect is well-documented and stems from polymer chain degradation, which reduces molecular weight and thereby viscosity ^38–40^. Specifically, glycosidic bonds of sodium alginate undergo hydrolytic cleavage, while cellulose derivatives (NaCMC, HEC) may be susceptible to oxidative degradation, potentially due to free radical formation of hydroperoxides. Notably, the extent of degradation can vary depending on the manufacturer and batch-related impurities, contributing to differences in viscosity loss ^41,42^. It is also important to note that the effects of sterilisation on gel viscosity can be reduced by the addition of electrolytes. For example, hydrogels made with NaCMC and HPMC showed an increase in viscosity, hardness, compressibility and adhesiveness in medium with higher ionic strength. The ionic stabilising effect was also observed with different grades of carbomers ^43^. Furthermore an older study suggests that propylene glycol (which is present at 25% in APOgel) may help stabilise alginate solutions by moderating temperature-dependent viscosity changes ^44^.

As defined in **Table 3**, the target viscosity profile of the sterile APOgel was within ±15% of the Nu-Gel benchmark product over a range of shear rates from 1 to 100 s^-1^, which represent shear stress values of common processes, including mixing and pouring ^40^. The criteria could largely be met by increasing the amount of HEC in the APOgel formulation to 2% w/w, resulting in a hydrogel with a higher viscosity pre-sterilisation (∼700 Pa*s) and a post-sterilisation viscosity profile roughly similar to the Nu-Gel product in both the initial and back shear rate sweep (i.e. within ±27% when comparing shear rates of 1, 10 and 100 s^-1^; **Figure 2A-C)**. Storage of the sterile APOgel in the syringes at 4°C over 90 days showed a trend towards increased viscosity with storage time **(Figure 2D)**.

**Figure 2:**
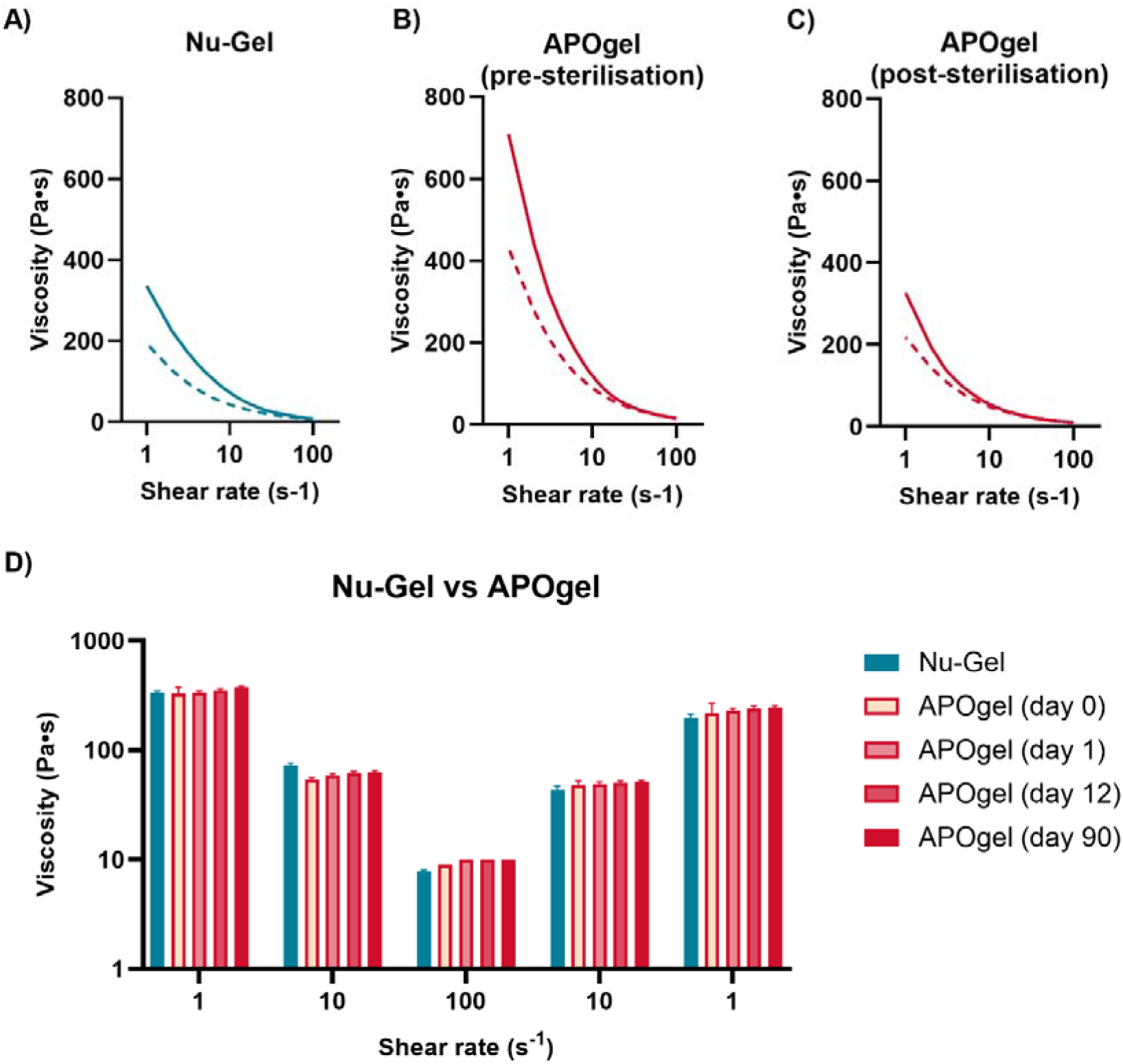
Dynamic viscosity values (Pa·s) across a shear rate sweep from 1 to 100 s^−1^for (A) Nu-Gel, (B) APOgel (pre-sterilisation), and (C) APOgel (post-sterilisation). Solid lines depict the first sweep, while dotted lines depict the back sweep. (D) Selected dynamic viscosity (Pa·s) values of Nu-Gel and sterile APOgel tested after storage at 4°C for 1, 12 and 90 days. Results represent the mean ± standard deviation of n=9 (three batches with three technical replicates per batch).

The reduction in APOgel viscosity following steam heat sterilisation did not induce noticeable amounts of degradation products as assessed by ATR-FTIR (**Figure 3A**). The ATR-FTIR spectra did not show measurable differences between Nu-Gel, APOgel (pre-sterilisation) and the sterile APOgel product (spectral similarity analysis is reported in the electronic supplementary information). To determine if sterilisation introduced acidic breakdown products, the pH values of the gels were pH following dilution with distilled water, since direct pH measurements of the gels proved challenging. Due to low conductivity, the pH of the distilled water showed a higher-than-expected pH of 9.54 ± 0.06, which, although not reflective of the true pH, is a common observation. Following dilution with distilled water, Nu-Gel and APOgel pre-sterilisation exhibited a similar pH-reduction (8.74 ± 0.20 and 8.73 ± 0.29, respectively) compared to water alone, while the pH of the diluted APOgel post-sterilisation was marginally but not significantly lower (8.54 ± 0.28; p=0.37), indicating that the sterilisation did not result in a meaningful increase in acidic functional groups. Based on the growing literature in the treatment of chronic, poorly healing wounds, evidence is mounting that more acidic pH topical preparations improve therapeutic outcomes via suppression of bacterial growth, facilitation of oxygenation, and stimulation of cellular signalling involved in wound healing ^45^. The pH of the vehicle gel may also impact the biological activity of APOSEC^TM^ components, which has not yet been investigated. Finally, the visual appearance of the APOgel formulations in the COP syringes were assessed pre-and post-sterilisation (**Figure 3B**). The sterilisation process increased the number of bubbles but reduced their overall size, and the colour appeared slightly yellowish. Overall, the investigated parameters suggest only very minor changes to the APOgel following sterilisation.

**Figure 3:**
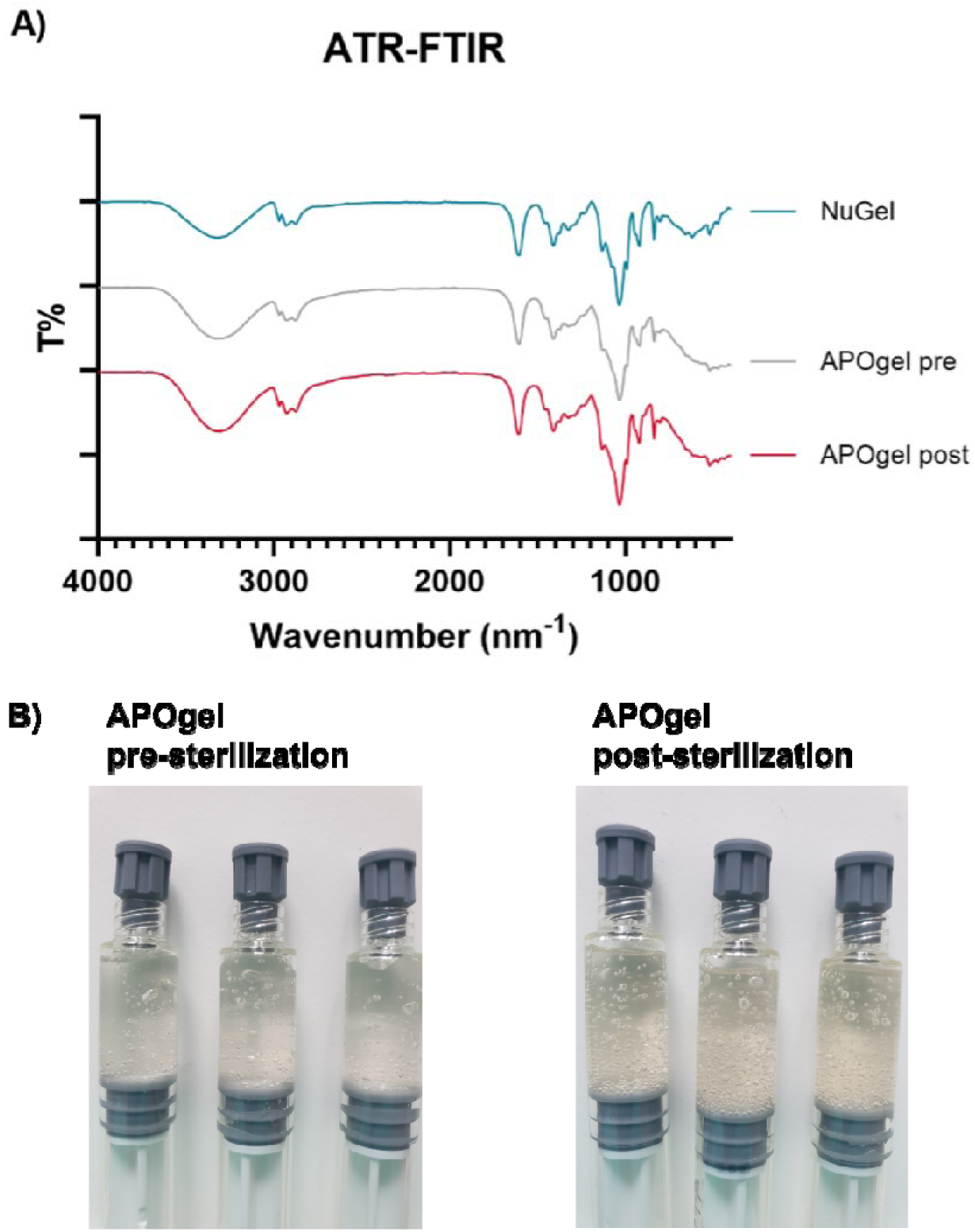
(A) ATR-FTIR spectra of Nu-Gel (blue) and APOgel (grey: pre-sterilisation; red: post-sterilisation). (B) Visual differences of APOgel pre- and post-sterilisation in COP syringes.

### 3.2 Pharmaceutical performance of the mixing/administration system

The proposed syringe-based mixing and dispensing system may be attractive for many putative applications but is also not well-characterised in the literature. To structure our current investigation, a list of CQAs and the corresponding target specifications for an envisioned APOSEC 2.0 dispensing system are listed in **Table 4**.

**Table 4:**
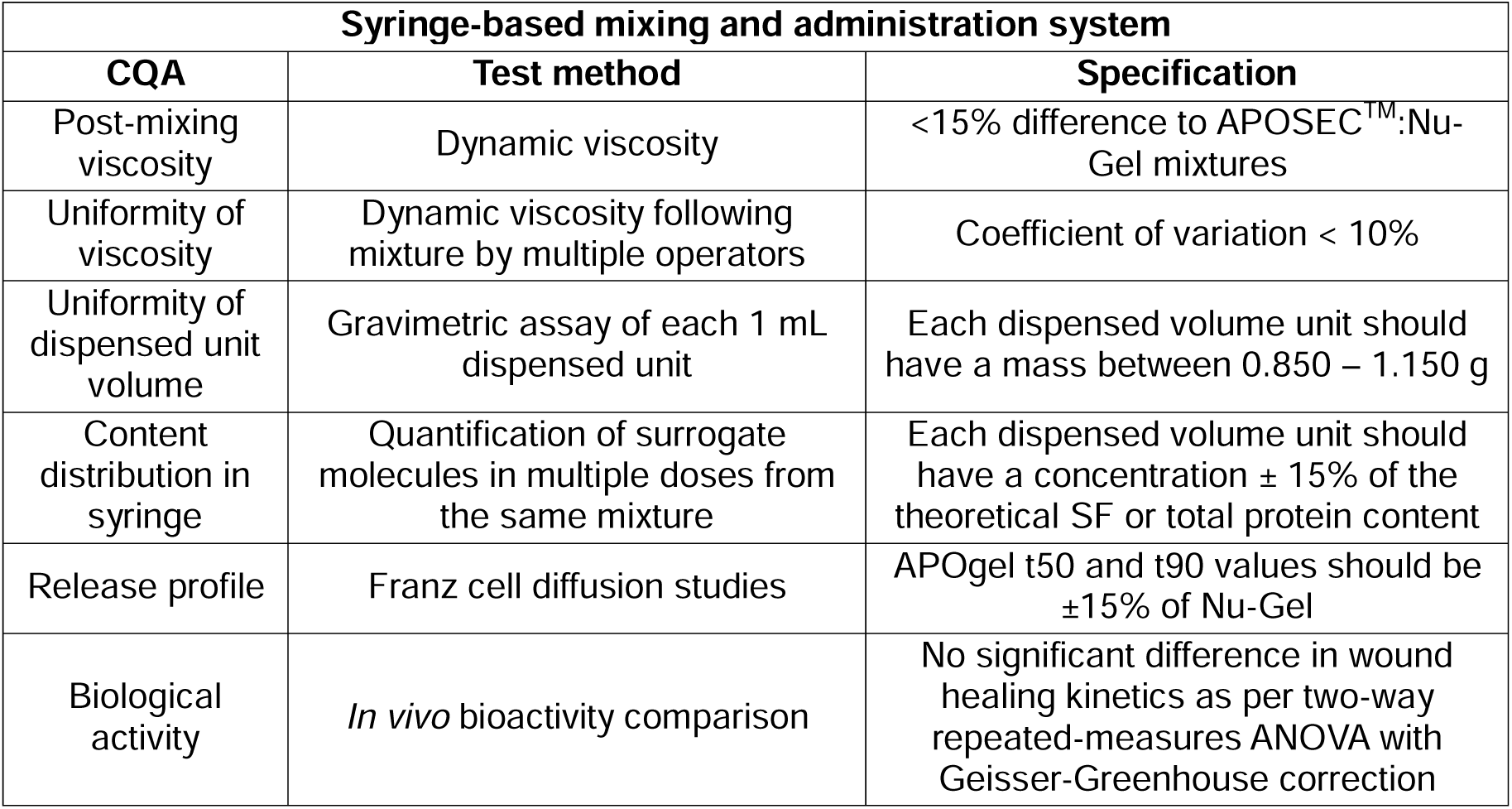
CQAs and target specifications for the putative APOSEC 2.0 syringe-mixing and dispensing system as illustrated in Figure 1.

#### 3.2.1 Mixture properties

To understand how the proposed syringe-mixing procedure will affect the properties of the APOSEC^TM^:APOgel mixture, pilot investigations were performed using AM2 medium instead of reconstituted APOSEC^TM^, which acted as a low-cost surrogate liquid with a similar viscosity. Preliminary studies found that optimal mixing of the liquid AM2 medium (1 g = 1mL) with 3 g APOgel occurred when the two components were combined and the mixture passed 20 times between the two syringes. The resulting viscosity of the AM2:NuGel mixture exhibited a viscosity reduction of 56% compared to the Nu-Gel alone (**Figure 4A**), when comparing at the initial shear rate of 1 s^-1^, reflective of the viscosity directly following administration from the syringe. The AM2:APOgel mixture (**Figure 4B**) exhibited a 67% viscosity reduction compared to the sterile APOgel and was also 27% less viscous than the AM2:Nu-Gel mixture. The lower viscosity of the AM2:APOgel mixture compared to Nu-Gel was outside the target specifications listed in **Table 4**. We further investigated whether the experience of operators resulted in different viscosities of AM2:APOgel mixtures, since preliminary studies indicated that participants tended to mix more carefully with less experience. It was hypothesised that with increased practice, participants might apply higher shear forces resulting in lower mixture viscosity ^46^. In our follow-on study, experience levels (no experience, some experience = 3–5 prior mixing operations, and experienced = >20 previous mixing operations) showed overall comparable viscosity to **Figure 4B** with a low variability (CV < 6%), suggesting no meaningful change in viscosity through repeated practice and compliance with our targeted specifications.

**Figure 4:**
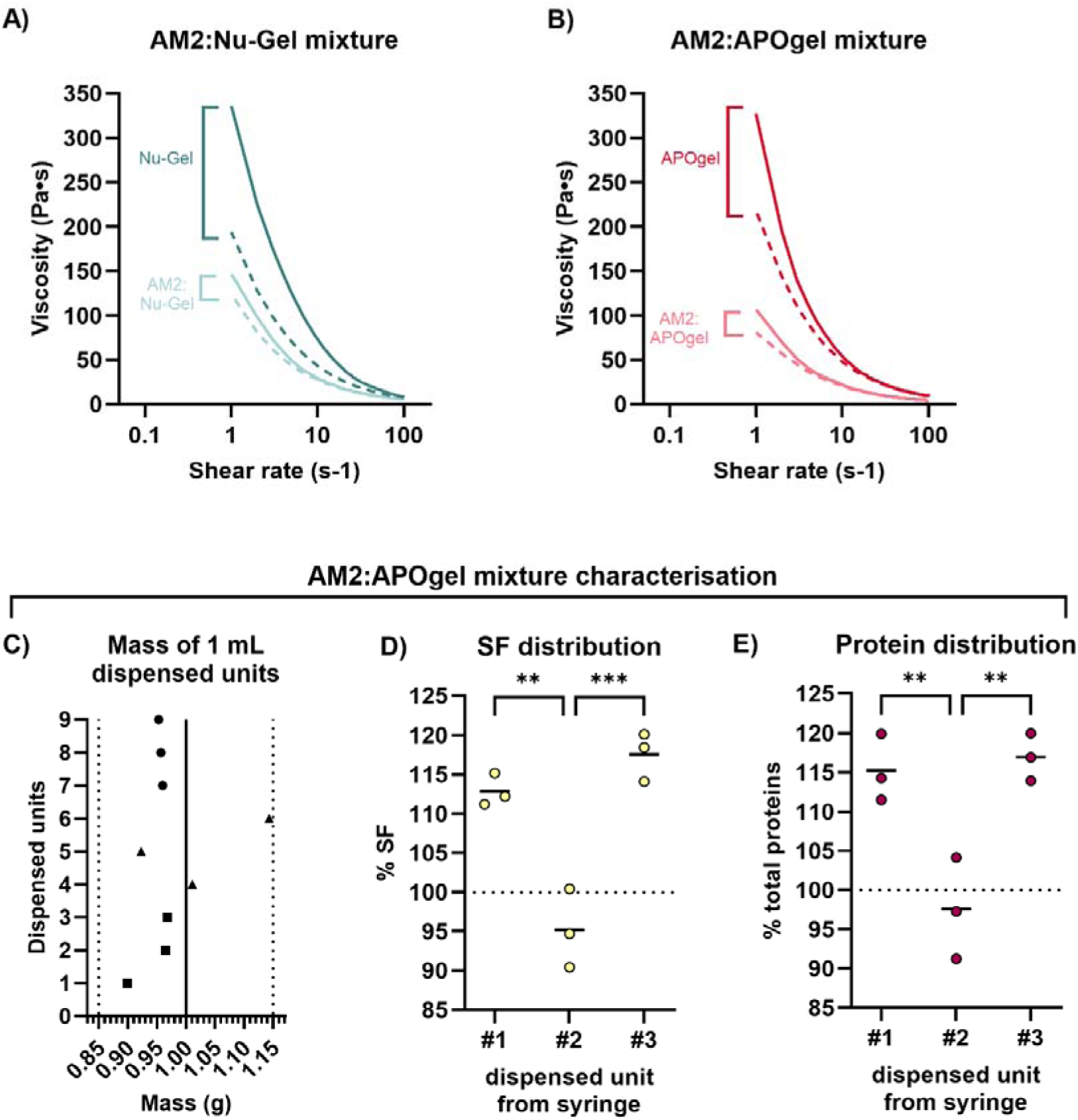
Dynamic viscosity values (Pa·s) across a shear rate sweep from 1 to 100 s^−1^for gels and AM2: gel mixtures (1:3). (A) Nu-Gel and (B) sterile APOgel. Viscosity profiles represent the mean of n=9 experiments using three different batches. (C) Mass values of nine dispensed unit volumes (1 mL) of AM2:APOgel across different operators. Each operator is indicated by a different symbol. Distribution of SF (D) and total proteins (E) in three sequential but spatially separated 0.1 g samples of AM2:APOgel mixture dispensed from three different syringes by a single experienced operator.

The consistency of dispensing 1 mL units of AM2:APOgel from the graduated syringe was assessed gravimetrically using three untrained volunteers (symbols indicate different volunteers; **Figure 4C**). It was assumed that a 1 mL unit volume dispensed from the graduated syringe would equate to 1 g. Eight of ten dispensed volumes were within ±10% of the expected mass, while all ten were within the ±15%. A trend toward slightly lower mass values was observed. To determine whether the syringe-mixing procedure achieved a homogenous distribution of AM2 components within each AM2:APOgel mixture, three samples of equal mass were dispensed sequentially from each syringe and the content of both a small molecule marker (SF) and total proteins was quantified. Surprisingly, the relative amounts of SF and total protein in samples #1 and #3 were consistently higher than sample #2, indicating a slight concentration gradient towards both ends of the syringe (**Figure 4D-E**). However, a mass balance was not performed, thus further variations in the gel may be present.

A recent study highlighted the challenges of creating a homogenous mixture via syringe-to-syringe mixing of viscous biomaterials, applying a similar method as described here ^47^. Although this study focussed on investigating the effects of different geometries of the mixing units on viscosity, they provided evidence that a discrepancy can occur between the front and the end of the syringe. The significance of analysing syringe concentration gradients is further highlighted by Xing *et al.*, where fluid dynamics within syringes were shown to cause uneven protein distribution ^48^. Although all fractions of the tested AM2:APOgel mixtures were within a ±25% range of the mean content and the majority within ±15%, this variability raises concerns for dose accuracy when only part of the syringe volume is administered. Further optimisation of the mixing procedure or alternative device geometries will be necessary to ensure more consistent distribution across the syringe barrel.

#### 3.2.2 *In vitro* release studies

Release studies of analytes from the AM2:gel mixtures were conducted using Franz-type diffusion cells ^49^, a standard *in vitro* method for simulating transdermal delivery through either skin tissue or, in the case of this study, synthetic membranes ^50^. To evaluate the release plus membrane diffusion rates of both small and large molecules from the mixture, SF (0.025% *w/v*) was added to the AM2 medium prior to mixing. Since AM2 medium also contains insulin and albumin, the total protein content release from the mixture was also analysed. A 6-h study (**Figure 5A,C**) was performed to assess initial release rates under high gradient, infinite dose (500 mg sample) conditions, while a 72-h study was performed to assess component release over a therapeutically relevant timeframe (i.e. a proposed dosing frequency of once every three days; **Figure 5B,D**). In the latter case, the medium exchange occurred only once every 24-h resulting in a lower concentration gradient and back-diffusion. A low viscosity control comprised of water, PPG and NaCl (in the same amounts as the gel formulations) was prepared and mixed with AM2 medium to highlight the effects of the mixture viscosity on the release rate of small and large molecular weight compounds.

**Figure 5:**
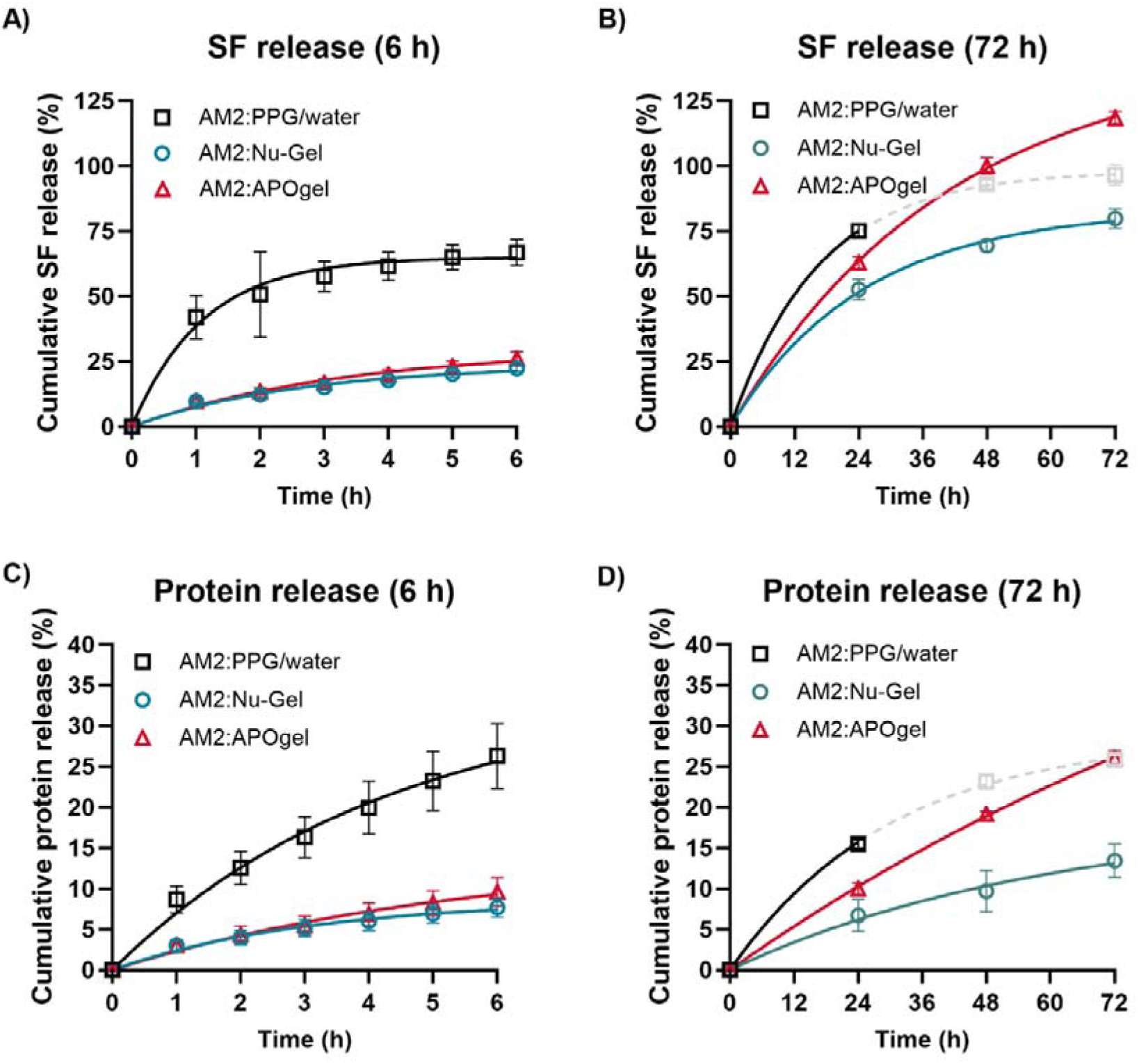
In vitro cumulative release plus membrane diffusion (%) profiles of SF and total proteins over 6 h (A, C; high gradient conditions) and 72 h (B, D; lower gradient conditions) from AM2:gel mixtures. A low viscosity control mixture (AM2:PPG/water) was included to highlight the effects of the gel component of the mixture on release profiles. Coloured lines represent fitting of the curves to Equation 1. The 48 and 72 h time points in the AM2:PPG/water controls are indicated in grey due to a loss of donor fluid at these time points and therefore a higher uncertainty in the data. Data points represent the mean ± standard deviation of n=3 batches.

Release plus membrane diffusion profiles under high gradient conditions (6-h study) showed more rapid diffusion of SF and protein from the low viscosity control sample compared to the gels (**Figure 5A,C**), verifying that the gels slow the release of both small molecules and proteins ^51^. In both cases, the calculated release rate constant *K*(*r*) of the AM2:PPG/water control was higher than for both gel mixtures (**Table 5**). Comparison of the SF and protein release from the AM2:gel mixtures revealed a slightly more rapid release when APOgel was used (**Figure 5A,C**), likely due to the lower viscosity compared to Nu-Gel mixtures (**Figure 4A-B**).

**Table 5:**
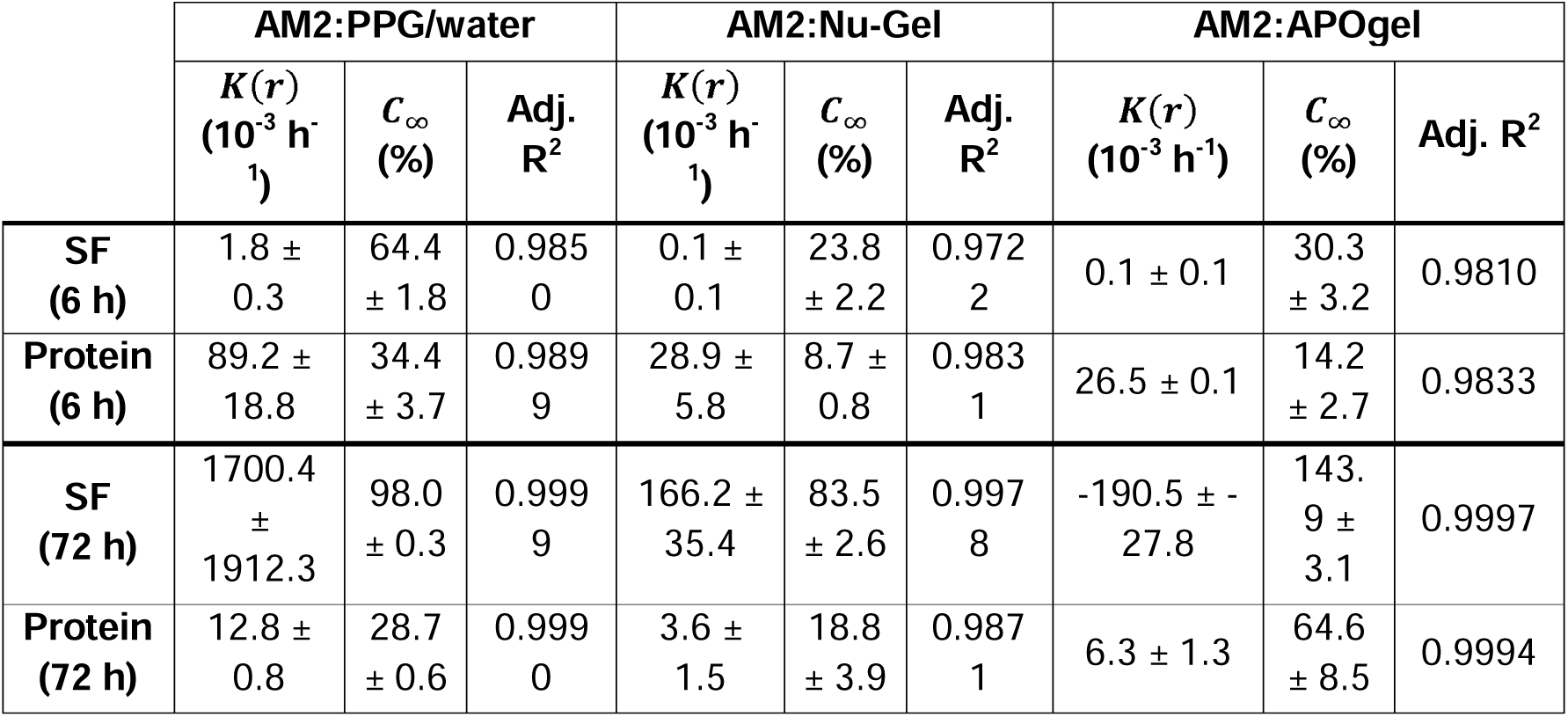
Release rate constant (K(r); 10^-3^ h^-1^) and predicted maximum release (C_∞_; %) of SF and proteins from samples mixed with AM2, obtained by fitting the release profiles to Equation 1.

The fitted release profiles of SF and proteins over 72 h (**Figure 5B,D**) revealed a more rapid and extensive release of SF and total proteins, despite the likely occurrence of back-diffusion. It should be noted that the profiles of the low viscosity controls are depicted in black only for the first 24 h, where a measurable amount of fluid was still present in the donor chamber. After this time, gravimetrical analysis of the changes to total mass of the system (**Figure 6**) indicated that the donor chamber of the AM2:PPG/water samples may have been depleted and therefore the concentrations of the later time points are shown in grey. The mass measurements presented in **Figure 6** indicate that the donor chambers of the AM2:gel mixtures did not deplete but rather increased in mass. The diffusion rate of SF and total proteins from AM2:APOgel showed even higher values over 72 h compared to diffusion from AM2:Nu-Gel mixtures. A rationale for these discrepancies can be drawn from formulation differences, including the impact that sterilisation likely had on the APOgel network structure, crosslinking density and viscosity, which would likely influence the release profiles differently^38^. The sustained and higher release observed for the AM2:APOgel mixture, may be of therapeutic benefit when extended drug availability is desired over a 72-h period. **Table 5** reports the comparative release rate constant and predicted maximum release values for all release profiles of **Figure 5**, according to **Equation 1**.

**Figure 6:**
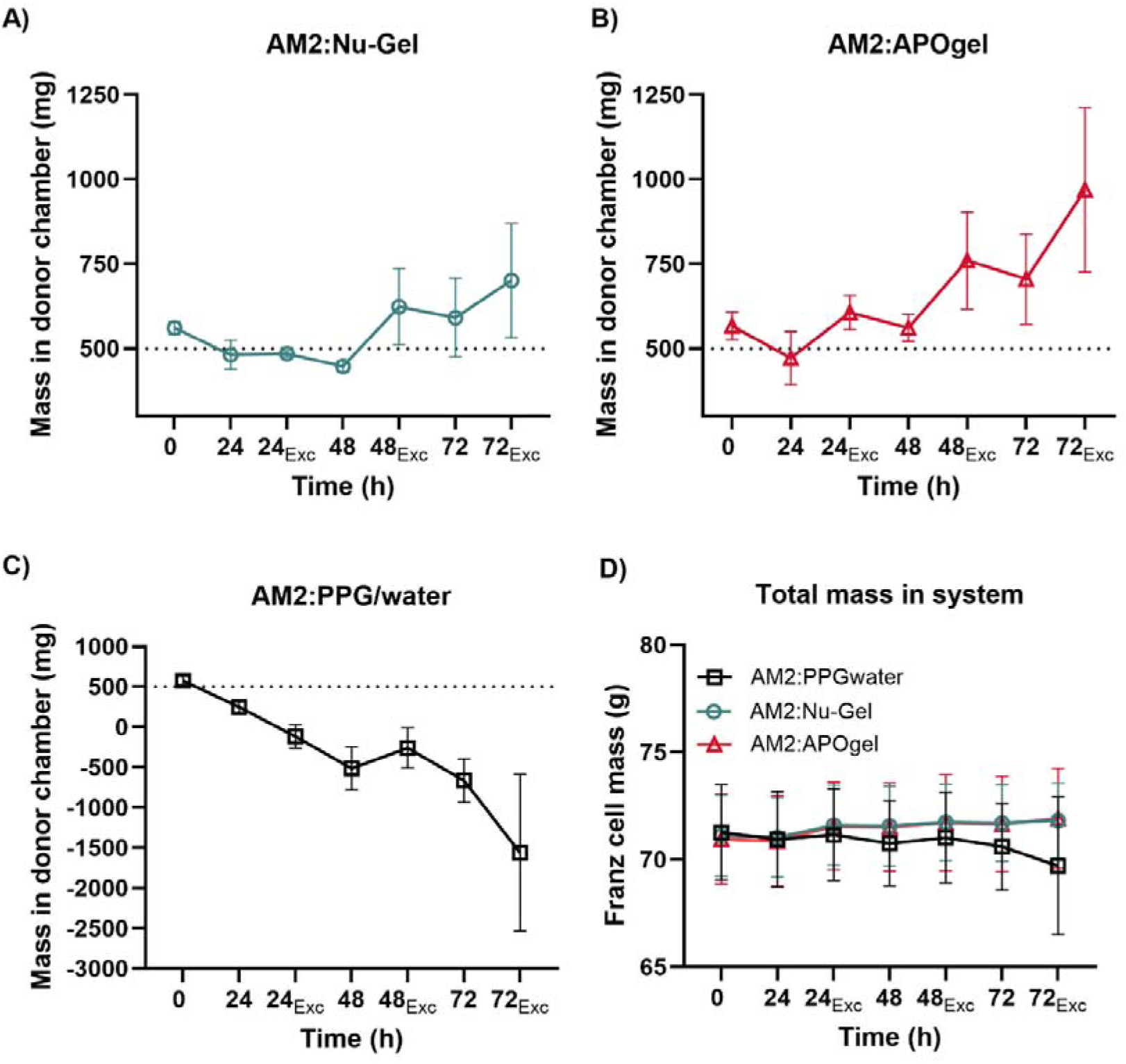
Change in sample mass within the donor chamber over 72 h relative to time point 0 h: (A) AM2:Nu-Gel, (B) AM2:APOgel, (C) AM2:PPG/water. (D) Changes to the total mass of the Franz cell at different time points of the experiment. The abbreviated subscript, Exc, indicates a medium exchange, i.e. the mass measured after removal of the entire volume of the acceptor chamber and addition of 2 mL buffer. Values represent the mean ± standard deviation of n=6 samples (two replicate experiments with three gel batches).

Tracking of the total Franz cell mass and calculation of the donor compartment mass at each time point before and after medium exchange in the 72-h study provided insights into concurrent evaporation in the system and swelling of the gels. (**Figure 6A-B**). In the first 48 h, the AM2:Nu-Gel samples exhibited mild losses due to evaporation and gel swelling was only observable in the final 24 h. The AM:APOgel mixtures, in contrast, showed evidence of substantial swelling behaviour at all time points, as seen in the increases in mass following medium exchange. The enhanced swelling of the AM2:APOgel system is another indication of a sterilisation-induced alteration of the APOgel structure and lower viscosity, which in a therapeutic context could promote both compound release and exudate uptake, effects which are beneficial for wound healing ^52,53^.

#### 3.2.3 *In vivo* wound healing kinetics following APOSEC^TM^ hydrogel treatment

To compare the *in vivo* wound-healing efficacy of APOSEC^TM^ incorporated into either Nu-Gel or APOgel, Balb/C mice were treated for ten days following full-thickness excisional wounding. A schematic overview of the experimental design is shown in **Figure 7A**. The respective APOSEC^TM^-gel mixtures were applied to the wound surface every second day, and the healing process was documented daily through standardized photography under identical conditions. Wound areas were quantified using ImageJ by manually tracing the wound borders and the percentual remaining wound area was calculated relative to the day of maximum wound expansion for each animal on each day. One animal in the Nu-Gel group (M4) had to be excluded from the analysis, as it had repetitively scratched and thereby mechanically enlarged its wound during the observation period, resulting in non-comparable wound measurements. The final dataset therefore included APOgel (n = 5) and Nu-Gel (n = 4) mice.

**Figure 7:**
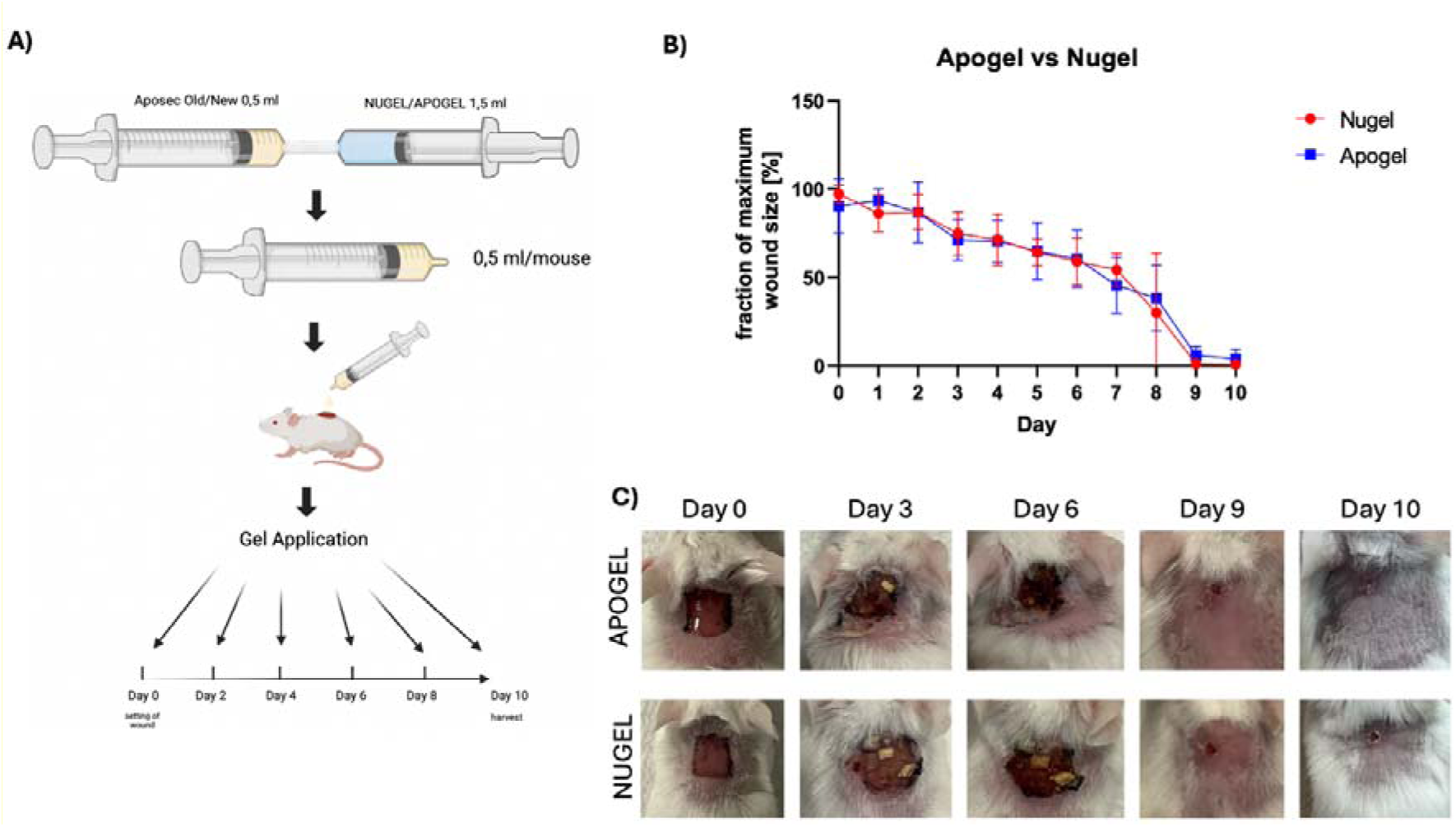
Comparative in vivo evaluation of APOSEC^TM^ hydrogel formulations. (A) Schematic of the in vivo wound healing experiment. Lyophilised APOSEC^TM^ was reconstituted in 0.9% NaCl, mixed with hydrogel (Nu-Gel or APOgel) and applied topically every second day for 10 days. (B) Wound closure kinetics of APOgel (blue) and Nu-Gel (red) expressed as fraction of maximum wound size (%). Data are mean ± SD (APOgel n = 5, Nu-Gel n = 4). Two-way repeated-measures ANOVA with Geisser–Greenhouse correction showed no significant difference between groups (treatment p = 0.94; interaction p = 0.83). (C) Representative wound images at days 0, 3, 6, 9 and 10 showing comparable healing progression in both treatment groups.

Statistical analysis was performed using two-way repeated-measures ANOVA with Geisser–Greenhouse correction in GraphPad Prism 9.0. The factors included time (within-subject) and treatment group (between-subject). As shown in **Figure 7B**, wound size significantly decreased over time in both groups (*p* < 0.0001), confirming progressive healing. No significant effect of treatment (*p* > 0.05) or interaction between time and treatment (*p >* 0.05) was detected, indicating that the healing kinetics were comparable between APOSEC^TM^:Nu-Gel - and APOSEC^TM^:APOgel -treated mice. Representative wound images from individual animals are displayed in **Figure 7C**, illustrating the similar macroscopic progression of wound closure in both treatment groups. Overall, these results demonstrate that the APOSEC^TM^:APOgel formulation performs equivalently to APOSEC^TM^:Nu-Gel in an in vivo wound healing assay, despite the differences in in vitro release profiles.

### 3.3 Interpretation of findings beyond APOSEC^TM^

Hydrogels are already established as versatile carriers with tuneable mechanics, including their capability to form injectable depots that localise and control the release of bioactive agents ranging from small molecules to therapeutic proteins and EVs ^54^. Integrating biologics and EVs into hydrogels is particularly attractive for regenerative medicine, as it can extend their bioactivity, improve stability, and overcome the rapid clearance that limits their efficacy *in vivo* ^55,56^. Yet, moving these systems toward clinical application requires overcoming technical challenges, including potentially slower release kinetics of macromolecules and EVs, possible effects of mixing on EV structure and function (which was not studied here) and achievement of a uniform distribution within the hydrogel matrix. This study attempts to fill some of these knowledge gaps whereby the development of an autoclavable hydrogel and a syringe-based mixing process showed some promising aspects.

## 4 Conclusion

This study demonstrates the investigation of a autoclavable hydrogel, APOgel, as a sterile matrix for topical administration of a biological product, using APOSEC^TM^ as an example system. The proposed gel formulation met several of the predefined CQAs, including sterility, pH, and chemical stability but also exhibited a slightly lower post-sterilisation viscosity compared the benchmark system, Nu-Gel. The performance of the closed syringe mixing system used to combine the gel and reconstituted liquid prior to administration revealed an uneven distribution of small-molecules and proteins in the syringe, an observation that highlights a limiting factor for homogenous dispensing from a single syringe. The lower viscosity of liquid:APOgel mixtures resulted in a more rapid release of both small molecule markers and proteins from the mixture, compared to the benchmark, which may have potential benefits for the overall therapeutic effect. However, in vivo wound-healing experiments demonstrated that APOSEC^TM^:APOgel formulation performed equivalently to the more viscous APOSEC^TM^:Nu-Gel system, demonstrating that the in vitro release profiles may not fully reflect in vivo conditions. Overall, these findings show that sterilised APOgel represents a viable hydrogel platform for topical administration to open wounds, while the proposed syringe mixing system could benefit from further optimisation.

## Acknowledgements

We thank our colleagues at the University of Vienna and the Medical University of Vienna for their valuable contributions. We also extend our gratitude to the colleagues at LKH-University Hospital Graz for their support and for providing access to their facilities. We thank H.P. Haselsteiner and Karl Fister, head of the CRISCAR Familienstiftung, for their faith in this private public partnership to augment basic and translational clinical research at the Medical University and University of Vienna.

## Author contributions

Dalia Hamid: conceptualization, writing - original draft, investigation (rheometry, FT-IR, gel production, uniformity studies, release study), formal analysis

Lisa Auer: conceptualization, investigation (animal experiments), writing – original draft, formal analysis

Sylwia Gazda-Miarecka: data curation, validation (documentation and quality assurance)

Stefan Mohr: resources, methodology (GMP gel production)

Victoria Klang: methodology

Tanja Pfleger: methodology

Melanie Salek: investigation (animal experiments), conceptualization

Hannes Kühtreiber: visualization, investigation (animal experiments), conceptualization

Nina Langoth-Fehringer: conceptualization, formal analysis (interpretation), validation (regulatory)

Michael Mildner: conceptualization, formal analysis, investigation, supervision

Clemens Aigner: resources, investigation, conceptualization

Dirk Sorgenfrey: conceptualization, formal analysis, writing – review and editing, validation

Jan H. Ankersmit: conceptualization, investigation, supervision, writing – review and editing, resources, methodology

Lea Ann Dailey: visualisation, methodology, supervision, conceptualization, formal analysis, writing – review and editing, resources, validation, investigation, project administration

Gianluca Bello: conceptualization, visualisation, methodology, investigation, data curation, supervision, writing – review and editing, formal analysis, validation, project administration, software,

## References

(1) Monaghan, M. G.; Borah, R.; Thomsen, C.; Browne, S. Thou Shall Not Heal: Overcoming the Non-Healing Behaviour of Diabetic Foot Ulcers by Engineering the Inflammatory Microenvironment. Advanced Drug Delivery Reviews 2023, 203, 115120. 10.1016/j.addr.2023.115120.

(2) Deng, H.; Li, B.; Shen, Q.; Zhang, C.; Kuang, L.; Chen, R.; Wang, S.; Ma, Z.; Li, G. Mechanisms of Diabetic Foot Ulceration: A Review. J Diabetes 2023, 15 (4), 299–312. 10.1111/1753-0407.13372.

(3) Hacker, S.; Mittermayr, R.; Nickl, S.; Haider, T.; Lebherz-Eichinger, D.; Beer, L.; Mitterbauer, A.; Leiss, H.; Zimmermann, M.; Schweiger, T.; Keibl, C.; Hofbauer, H.; Gabriel, C.; Pavone-Gyöngyösi, M.; Redl, H.; Tschachler, E.; Mildner, M.; Ankersmit, H. J. Paracrine Factors from Irradiated Peripheral Blood Mononuclear Cells Improve Skin Regeneration and Angiogenesis in a Porcine Burn Model. Sci Rep 2016, 6 (1), 25168. 10.1038/srep25168.

(4) Wagner, T.; Traxler, D.; Simader, E.; Beer, L.; Narzt, M.-S.; Gruber, F.; Madlener, S.; Laggner, M.; Erb, M.; Vorstandlechner, V.; Gugerell, A.; Radtke, C.; Gnecchi, M.; Peterbauer, A.; Gschwandtner, M.; Tschachler, E.; Keibl, C.; Slezak, P.; Ankersmit, H. J.; Mildner, M. Different Pro-Angiogenic Potential of γ-Irradiated PBMC-Derived Secretome and Its Subfractions. Sci Rep 2018, 8, 18016. 10.1038/s41598-018-36928-6.

(5) Beer, L.; Mildner, M.; Gyöngyösi, M.; Ankersmit, H. J. Peripheral Blood Mononuclear Cell Secretome for Tissue Repair. Apoptosis 2016, 21 (12), 1336–1353. 10.1007/s10495-016-1292-8.

(6) Beer, L.; Zimmermann, M.; Mitterbauer, A.; Ellinger, A.; Gruber, F.; Narzt, M.-S.; Zellner, M.; Gyöngyösi, M.; Madlener, S.; Simader, E.; Gabriel, C.; Mildner, M.; Ankersmit, H. J. Analysis of the Secretome of Apoptotic Peripheral Blood Mononuclear Cells: Impact of Released Proteins and Exosomes for Tissue Regeneration. Sci Rep 2015, 5 (1), 16662. 10.1038/srep16662.

(7) Laggner, M.; Gugerell, A.; Copic, D.; Jeitler, M.; Springer, M.; Peterbauer, A.; Kremslehner, C.; Filzwieser-Narzt, M.; Gruber, F.; Madlener, S.; Erb, M.; Widder, J.; Lechner, W.; Georg, D.; Mildner, M.; Ankersmit, H. J. Comparing the Efficacy of γ- and Electron-Irradiation of PBMCs to Promote Secretion of Paracrine, Regenerative Factors. Mol Ther Methods Clin Dev 2021, 21, 14–27. 10.1016/j.omtm.2021.02.016.

(8) Beer, L.; Seemann, R.; Ristl, R.; Ellinger, A.; Kasiri, M. M.; Mitterbauer, A.; Zimmermann, M.; Gabriel, C.; Gyöngyösi, M.; Klepetko, W.; Mildner, M.; Ankersmit, H. J. High Dose Ionizing Radiation Regulates Micro RNA and Gene Expression Changes in Human Peripheral Blood Mononuclear Cells. BMC Genomics 2014, 15 (1), 814. 10.1186/1471-2164-15-814.

(9) Laggner, M.; Gugerell, A.; Bachmann, C.; Hofbauer, H.; Vorstandlechner, V.; Seibold, M.; Gouya Lechner, G.; Peterbauer, A.; Madlener, S.; Demyanets, S.; Sorgenfrey, D.; Ostler, T.; Erb, M.; Mildner, M.; Ankersmit, H. J. Reproducibility of GMP-Compliant Production of Therapeutic Stressed Peripheral Blood Mononuclear Cell-Derived Secretomes, a Novel Class of Biological Medicinal Products. Stem Cell Research & Therapy 2020, 11 (1), 9. 10.1186/s13287-019-1524-2.

(10) Wuschko, S.; Gugerell, A.; Chabicovsky, M.; Hofbauer, H.; Laggner, M.; Erb, M.; Ostler, T.; Peterbauer, A.; Suessner, S.; Demyanets, S.; Leuschner, J.; Moser, B.; Mildner, M.; Ankersmit, H. J. Toxicological Testing of Allogeneic Secretome Derived from Peripheral Mononuclear Cells (APOSEC): A Novel Cell-Free Therapeutic Agent in Skin Disease. Sci Rep 2019, 9 (1), 5598. 10.1038/s41598-019-42057-5.

(11) Gugerell, A.; Sorgenfrey, D.; Laggner, M.; Raimann, J.; Peterbauer, A.; Bormann, D.; Suessner, S.; Gabriel, C.; Moser, B.; Ostler, T.; Mildner, M.; Ankersmit, H. J. Viral Safety of APOSECTM: A Novel Peripheral Blood Mononuclear Cell Derived-Biological for Regenerative Medicine. Blood Transfusion 2020, 30–39. 10.2450/2019.0249-18.

(12) Simader, E.; Traxler, D.; Kasiri, M. M.; Hofbauer, H.; Wolzt, M.; Glogner, C.; Storka, A.; Mildner, M.; Gouya, G.; Geusau, A.; Fuchs, C.; Eder, C.; Graf, A.; Schaden, M.; Golabi, B.; Aretin, M.-B.; Suessner, S.; Gabriel, C.; Klepetko, W.; Tschachler, E.; Ankersmit, H. J. Safety and Tolerability of Topically Administered Autologous, Apoptotic PBMC Secretome (APOSEC) in Dermal Wounds: A Randomized Phase 1 Trial (MARSYAS I ). Scientific Reports 2017, 7, 6216. 10.1038/s41598-017-06223-x.

(13) MARSYAS II Synopsis of Clinical Study Report – Aposcience AG. https://www.aposcience.at/news/marsyas-ii-synopsis-of-clinical-study-report/ (accessed 2026-01-08).

(14) Ma, T.; Xu, G.; Gao, T.; Zhao, G.; Huang, G.; Shi, J.; Chen, J.; Song, J.; Xia, J.; Ma, X. Engineered Exosomes with ATF5-Modified mRNA Loaded in Injectable Thermogels Alleviate Osteoarthritis by Targeting the Mitochondrial Unfolded Protein Response. ACS Appl. Mater. Interfaces 2024, 16 (17), 21383–21399. 10.1021/acsami.3c17209.

(15) Hwang, H. S.; Lee, C.-S. Exosome-Integrated Hydrogels for Bone Tissue Engineering. Gels 2024, 10 (12), 762. 10.3390/gels10120762.

(16) Zhang, Y.; Yan, W.; Wu, L.; Yu, Z.; Quan, Y.; Xie, X. Different Exosomes Are Loaded in Hydrogels for the Application in the Field of Tissue Repair. Front. Bioeng. Biotechnol. 2025, 13, 1545636. 10.3389/fbioe.2025.1545636.

(17) Fu, Y.; Qiu, Z.; Cao, Y.; Jiang, M.; Cui, X. Hydrogel–Exosome Complexes: A Novel Strategy for Cardiovascular Regeneration. Nanoscale 2025, 17 (24), 14477–14490. 10.1039/D5NR00892A.

(18) Knopp, J. L.; Holder-Pearson, L.; Chase, J. G. Insulin Units and Conversion Factors: A Story of Truth, Boots, and Faster Half-Truths. J Diabetes Sci Technol 2018, 13 (3), 597–600. 10.1177/1932296818805074.

(19) Larmené-Beld, K.; Wijnsma, R.; Kuiper, A.; Van Berkel, S.; Robben, H.; Taxis, K.; Frijlink, H. Science- and Risk-Based Strategy to Qualify Prefillable Autoclavable Syringes as Primary Packaging Material. Eur J Hosp Pharm 2022, 29 (5), 248–254. 10.1136/ejhpharm-2020-002333.

(20) Hardy, Craig Julian; Findlay, Charlotte Maria. Sterile Gel Compositions for Wound Treatment. EP0693292A1, 01 1996.

(21) S. A. Bento, C.; Gaspar, M. C.; Coimbra, P.; De Sousa, H. C.; E. M. Braga, M. A Review of Conventional and Emerging Technologies for Hydrogels Sterilization. International Journal of Pharmaceutics 2023, 634, 122671. 10.1016/j.ijpharm.2023.122671.

(22) Abraham, J. International Conference On Harmonisation Of Technical Requirements For Registration Of Pharmaceuticals For Human Use. In Handbook of Transnational Economic Governance Regimes; Tietje, C., Brouder, A., Eds.; Brill | Nijhoff, 2010; pp 1041–1053. 10.1163/ej.9789004163300.i-1081.897.

(23) De La Guardia, C.; Virno, A.; Musumeci, M.; Bernardin, A.; Silberberg, M. B. Rheologic and Physicochemical Characteristics of Hyaluronic Acid Fillers: Overview and Relationship to Product Performance. Facial Plast Surg 2022, 38 (02), 116–123. 10.1055/s-0041-1741560.

(24) Ayouch, I.; Kassem, I.; Kassab, Z.; Barrak, I.; Barhoun, A.; Jacquemin, J.; Draoui, K.; Achaby, M. E. Crosslinked Carboxymethyl Cellulose-Hydroxyethyl Cellulose Hydrogel Films for Adsorption of Cadmium and Methylene Blue from Aqueous Solutions. Surfaces and Interfaces 2021, 24, 101124. 10.1016/j.surfin.2021.101124.

(25) Phan, K. S.; Nguyen, T. M.; To, X. T.; Le, T. T. H.; Nguyen, T. T.; Pham, K. D.; Hoang, P. H.; Dong, T. N.; Dang, D. K.; Phan, T. H. T.; Mai, T. T. T.; Ha, P. T. Allium sativum@AgNPs and Phyllanthus urinaria@AgNPs: A Comparative Analysis for Antibacterial Application. RSC Adv 12 (55), 35730–35743. 10.1039/d2ra06847h.

(26) European Directorate for the Quality of Medicines & HealthCare (EDQM). European Pharmacopoeia (Ph. Eur.), 11th Edition: 2.6.1 Sterility. http://www.uspbpep.com/ep60/2.6.%201.%20sterility%2020601e.pdf (accessed 2025-09-30).

(27) Thermo Fisher Scientific. Pierce^TM^ Detergent Compatible Bradford Assay Kit User Guide; MAN0016018; Thermo Fisher Scientific, 2016. https://assets.thermofisher.com/TFS-Assets/LSG/manuals/23246_23246S_deter_compat_bradford_UG.pdf (accessed 2025-05-27).

(28) Bernik, D. L.; Zubiri, D.; Monge, M. E.; Negri, R. M. New Kinetic Model of Drug Release from Swollen Gels under Non-Sink Conditions. Colloids and Surfaces A: Physicochemical and Engineering Aspects 2006, 273 (1), 165–173. 10.1016/j.colsurfa.2005.08.018.

(29) Siepmann, J.; Siepmann, F. Sink Conditions Do Not Guarantee the Absence of Saturation Effects. International Journal of Pharmaceutics 2020, 577, 119009. 10.1016/j.ijpharm.2019.119009.

(30) Troiano, G.; Nolan, J.; Parsons, D.; Van Geen Hoven, C.; Zale, S. A Quality by Design Approach to Developing and Manufacturing Polymeric Nanoparticle Drug Products. AAPS J 2016, 18 (6), 1354–1365. 10.1208/s12248-016-9969-z.

(31) Dailey, L. A. Pharmaceutical Quality by Design in Academic Nanomedicine Research: Stifling Innovation or Creativity through Constraint? Journal of Interdisciplinary Nanomedicine 2018, 3 (4), 175–182. 10.1002/jin2.52.

(32) European Medicines Agency. Guideline on Quality and Equivalence of Locally Applied Locally Acting Cutaneous Products; EMA/CHMP/703337/2018; 2018. https://www.ema.europa.eu/en/documents/scientific-guideline/guideline-quality-equivalence-locally-applied-locally-acting-cutaneous-products_en.pdf (accessed 2025-08-10).

(33) European Medicines Agency (EMA). Guideline on the Sterilisation of the Medicinal Product, Active Substance, Excipient and Primary Container, 2019. https://www.ema.europa.eu/en/documents/scientific-guideline/guideline-sterilisation-medicinal-product-active-substance-excipient-and-primary-container_en.pdf.

(34) Dai, Z.; Ronholm, J.; Tian, Y.; Sethi, B.; Cao, X. Sterilization Techniques for Biodegradable Scaffolds in Tissue Engineering Applications. J Tissue Eng 2016, 7, 2041731416648810. 10.1177/2041731416648810.

(35) Galante, R.; Pinto, T. J. A.; Colaço, R.; Serro, A. P. Sterilization of Hydrogels for Biomedical Applications: A Review. J Biomed Mater Res 2018, 106 (6), 2472–2492. 10.1002/jbm.b.34048.

(36) Beard, M. C.; Cobb, L. H.; Grant, C. S.; Varadarajan, A.; Henry, T.; Swanson, E. A.; Kundu, S.; Priddy, L. B. Autoclaving of Poloxamer 407 Hydrogel and Its Use as a Drug Delivery Vehicle. J Biomed Mater Res 2021, 109 (3), 338–347. 10.1002/jbm.b.34703.

(37) Ahmed, E. M. Hydrogel: Preparation, Characterization, and Applications: A Review. J Adv Res 2015, 6 (2), 105–121. 10.1016/j.jare.2013.07.006.

(38) Stoppel, W. L.; White, J. C.; Horava, S. D.; Henry, A. C.; Roberts, S. C.; Bhatia, S. R. Terminal Sterilization of Alginate Hydrogels: Efficacy and Impact on Mechanical Properties. J Biomed Mater Res 2014, 102 (4), 877–884. 10.1002/jbm.b.33070.

(39) Alipour, S.; Negahban, N.; Ahmadi, F.; Parhizkar, E. The Effects of Moist Heat Sterilization Process on Rheological Properties of Hydrophilic Gels Containing Drug Model. Trends in Pharmaceutical Sciences 2022, 8 (3). 10.30476/tips.2022.94633.1138.

(40) Ferreira, I.; Marques, A. C.; Costa, P. C.; Amaral, M. H. Effects of Steam Sterilization on the Properties of Stimuli-Responsive Polymer-Based Hydrogels. Gels 2023, 9 (5), 385. 10.3390/gels9050385.

(41) Bakhrushina, E. O.; Afonina, A. M.; Mikhel, I. B.; Demina, N. B.; Plakhotnaya, O. N.; Belyatskaya, A. V.; Krasnyuk, I. I.; Krasnyuk, I. I. Role of Sterilization on In Situ Gel-Forming Polymer Stability. Polymers 2024, 16 (20), 2943. 10.3390/polym16202943.

(42) NatrosolTM 250 hydroxyethylcellulose (HEC). https://www.ashland.com/file_source/Ashland/links/PHA18-101_Natrosol_250_HEC_Formulating_elegant_liquid_and_semisolid_%20drug_products.pdf.

(43) Tichý, E.; Murányi, A.; Pšenková, J. The Effects of Moist Heat Sterilization Process and the Presence of Electrolytes on Rheological and Textural Properties of Hydrophilic Dispersions of Polymers-Hydrogels. Adv Polym Technol 2016, 35 (2), 198–207. 10.1002/adv.21543.

(44) Schwarz, T. W.; Levy, G. Viscosity Changes of Sodium Alginate Solutions after Freezing and Thawing**University of California School of Pharmacy, San Francisco. Journal of the American Pharmaceutical Association (Scientific ed.) 1957, 46 (9), 562–563. 10.1002/jps.3030460914.

(45) Sim, P.; Strudwick, X. L.; Song, Y.; Cowin, A. J.; Garg, S. Influence of Acidic pH on Wound Healing In Vivo: A Novel Perspective for Wound Treatment. IJMS 2022, 23 (21), 13655. 10.3390/ijms232113655.

(46) Uman, S.; Dhand, A.; Burdick, J. A. Recent Advances in Shear-thinning and Self-healing Hydrogels for Biomedical Applications. J of Applied Polymer Sci 2020, 137 (25), 48668. 10.1002/app.48668.

(47) Dani, S.; Ahlfeld, T.; Albrecht, F.; Duin, S.; Kluger, P.; Lode, A.; Gelinsky, M. Homogeneous and Reproducible Mixing of Highly Viscous Biomaterial Inks and Cell Suspensions to Create Bioinks. Gels 2021, 7 (4), 227. 10.3390/gels7040227.

(48) Xing, L.; Li, Y.; Li, T. Local Concentrating, Not Shear Stress, That May Lead to Possible Instability of Protein Molecules During Syringe Injection: A Fluid Dynamic Study with Two-Phase Flow Model. PDA Journal of Pharmaceutical Science and Technology 2019, 73 (3), 260–275. 10.5731/pdajpst.2018.009357.

(49) Franz, T. J. Percutaneous Absorption. On the Relevance of in Vitro Data. Journal of Investigative Dermatology 1975, 64 (3), 190–195. 10.1111/1523-1747.ep12533356.

(50) Ng, S.-F.; Rouse, J. J.; Sanderson, F. D.; Meidan, V.; Eccleston, G. M. Validation of a Static Franz Diffusion Cell System for In Vitro Permeation Studies. AAPS PharmSciTech 2010, 11 (3), 1432–1441. 10.1208/s12249-010-9522-9.

(51) Owh, C.; Ow, V.; Lin, Q.; Wong, J. H. M.; Ho, D.; Loh, X. J.; Xue, K. Bottom-up Design of Hydrogels for Programmable Drug Release. Biomaterials Advances 2022, 141, 213100. 10.1016/j.bioadv.2022.213100.

(52) Dumville, J. C.; O’Meara, S.; Deshpande, S.; Speak, K. Alginate Dressings for Healing Diabetic Foot Ulcers. Cochrane Database of Systematic Reviews 2013, 2015 (3). 10.1002/14651858.CD009110.pub3.

(53) A Randomized Controlled Trial: Effects of Alginate Gel in the Treatment of Diabetic Foot Ulcers. J Surg 2025, 10 (6). 10.29011/2575-9760.011316.

(54) Bhatta, R.; Han, J.; Liu, Y.; Bo, Y.; Wang, Y.; Nguyen, D.; Chen, Q.; Wang, H. Injectable Extracellular Vesicle Hydrogels with Tunable Viscoelasticity for Depot Vaccine. Nat Commun 2025, 16 (1), 3781. 10.1038/s41467-025-59278-0.

(55) Ju, Y.; Hu, Y.; Yang, P.; Xie, X.; Fang, B. Extracellular Vesicle-Loaded Hydrogels for Tissue Repair and Regeneration. Materials Today Bio 2023, 18, 100522. 10.1016/j.mtbio.2022.100522.

(56) Zheng, Y.; Pan, C.; Xu, P.; Liu, K. Hydrogel-Mediated Extracellular Vesicles for Enhanced Wound Healing: The Latest Progress, and Their Prospects for 3D Bioprinting. J Nanobiotechnol 2024, 22 (1), 57. 10.1186/s12951-024-02315-9.

