## Supplementary Info for "Investigation of sterile hydrogels as topical vehicles for APOSEC^TM^, a stressed peripheral blood mononuclear cell secretome for the treatment of poorly healing wounds"

### Figure S1 : Manufacturing process of APOSEC™

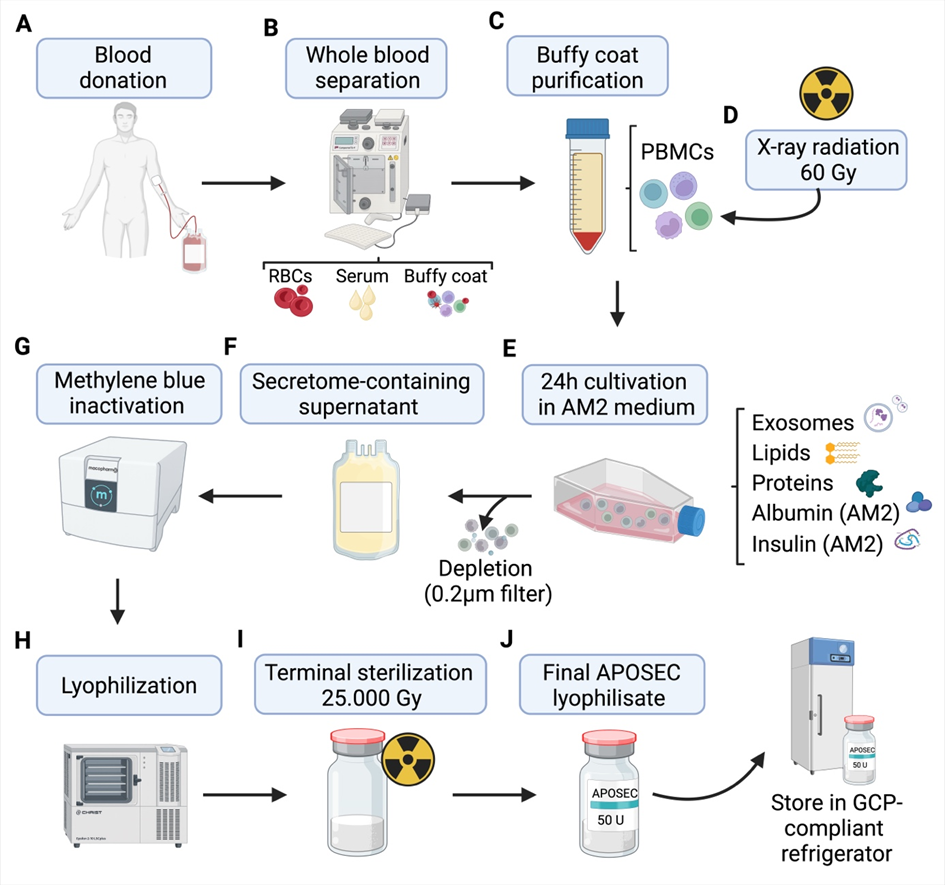

(A) Whole blood is collected from multiple healthy donors. (B) Using a Compomat G4 system (Fresenius Kabi), the blood is fractionated into approximately 200–250 mL serum, 200–250 mL packed red blood cells, and ~30 mL buffy coat per 500 mL donation. (C) The buffy coat fraction is centrifuged again to remove residual red blood cells, yielding a purified cell fraction. (D) Peripheral blood mononuclear cells (PBMCs) within the purified buffy coat are irradiated with 60 Gy X-rays. (E) Irradiated PBMCs are cultured for 24 h in AM2 medium, resulting in the release of exosomes, lipids, and proteins; insulin and albumin are present as integral components of the medium. (F) After depletion of cells and debris by filtration through a 0.2 µm filter, the secretome-containing supernatant is transferred into sterile storage bags as intermediate product. (G) Methylene blue inactivation is performed using the Macopharma system. (H) The product is subsequently lyophilized in a freeze-dryer (Lyophilisator, Martin Christ, Epsilon 2-10 LSCplus). (I) The lyophilizate is aliquoted into sterile vials (50 units APOSEC per vial) and subjected to terminal sterilization with 25,000 Gy. (J) The final APOSEC lyophilisate is stored in a GCP-compliant refrigerator until clinical use.

### Figure S2 : Preparation and administration of APOSEC™ TPP for the MARSYAS I/II trial

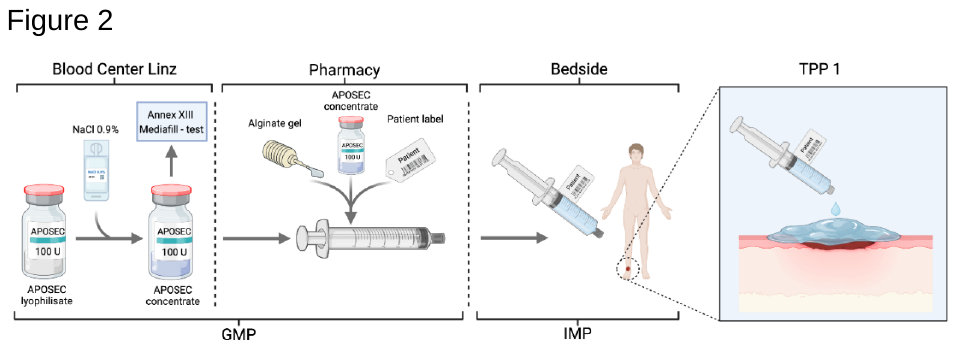

The frozen APOSEC^TM^ concentrate was distributed to the local pharmacy adjacent to the study site. The APOSEC concentrate was manually blended with the NUGEL alginate gel and filled into graduated syringe. The alginate gel - APOSEC mixture was distributed the study physician and applied to the wound of the randomized study patient.

#### ATR-FTIR similarity calculations

Similarity between different FTIR absorbance spectra was quantified using similarity measures with scores normalised to a range of 0–999, where 999 indicates identical spectra and values below 900 indicate not similar spectra. The calculations of 4 similarity measures are carried out with the Python (v3.13) code provided according to the theory reported in the study by Varmuza ^1^. A reference spectrum (in this case NUgel) is used and then compared to the other spectra (APOgel pre and post sterilization). The similarity measures focus on specific spectra aspects summarized in this table:

| **Measure** | **Focus** | **Score >900 meaning** |
| --- | --- | --- |
| COR | Pattern similarity | Peaks and valleys match |
| MAD | Average difference at each point | Spectra are close throughout the whole interval |
| MSD | Emphasizes large differences | No mismatches between spectra |
| DPN | Overall shape/intensity | Spectra point in same direction |

The FTIR spectra of APOgel pre and post sterilization did not show measurable differences to Nu-Gel, except for the DPN similarity of APOgel pre sterilization which differ only for 1 point. This could be to a normal variability between spectra and is not affecting the overall outcome.

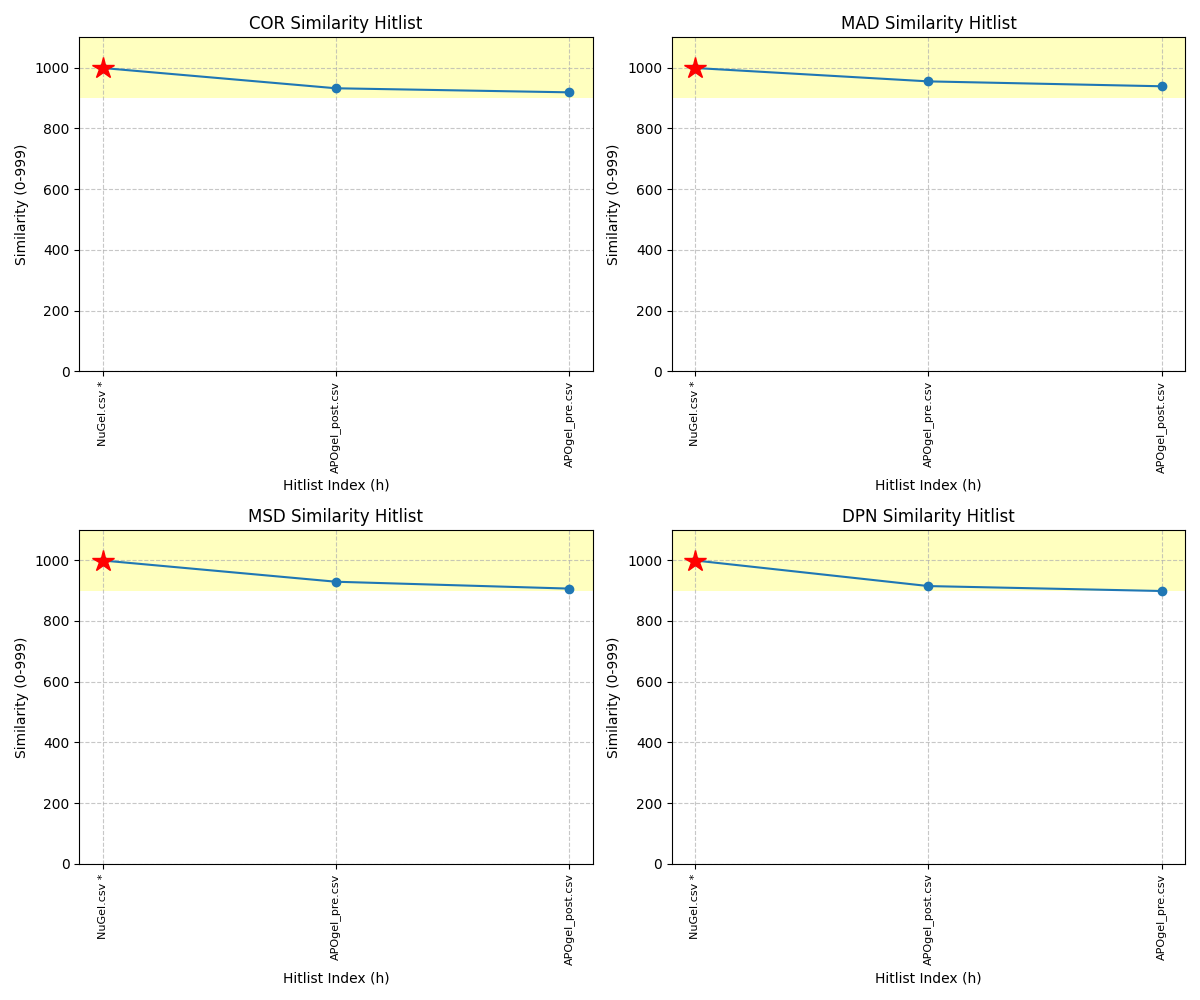

| Spectrum | COR | COR Similarity | MAD | MAD Similarity | MSD | MSD Similarity | DPN | DPN Similarity |
| --- | --- | --- | --- | --- | --- | --- | --- | --- |
| NuGel * | 999 | Similar | 999 | Similar | 999 | Similar | 999 | Similar |
| APOgel_post | 932 | Similar | 955 | Similar | 929 | Similar | 915 | Similar |
| APOgel_pre | 919 | Similar | 939 | Similar | 906 | Similar | 899 | Not Similar |

(1)	Varmuza, K.; Karlovits, M.; Demuth, W. Spectral Similarity versus Structural Similarity: Infrared Spectroscopy. *Analytica Chimica Acta* **2003**, *490* (1), 313–324. https://doi.org/10.1016/S0003-2670(03)00668-8.
